# Personality, Subjective Well-Being, and the Serotonin 1a Receptor Gene in Common Marmosets (*Callithrix jacchus*)

**DOI:** 10.1101/2020.04.30.069773

**Authors:** Alexander Weiss, Chihiro Yokoyama, Takuya Hayashi, Miho Inoue-Murayama

**Author notes:** **Author Note** We have no known conflicts of interest to declare. We are grateful to Hiromi Kobayashi for technical support and to Mr. Akihiro Kawasaki, Mr. Takashi Fukuoka, and Ms. Chiho Takeda for rating the animals. AW thanks Kyoto University for hosting him as a Visiting Professor at the Wildlife Research Center. Corresponding author (AW).

## Abstract

Studies of personality traits in common marmosets (*Callithrix jacchus*) indicate that there are five or six constructs—sociability, dominance, neuroticism, openness, and two related to conscientiousness—that define personality in common marmosets. The present study attempted to determine whether our earlier study of laboratory-housed individuals only yielded three domains—Dominance, Sociability, and Neuroticism—because of a low amount of between-subjects variance. We therefore increased our sample size from 77 to 128. In addition, we ascertained the reliability and validity of ratings and whether polymorphisms related to the serotonin 1a receptor were associated with personality. We found Sociability, Dominance, and Negative Affect factors that resembled three domains found in previous studies, including ours. We also found an Openness and Impulsiveness factor, the latter of which bore some resemblance to Conscientiousness, and two higher-order factors, Pro-sociality and Boldness. Further analyses could not exclude the possibility that Pro-sociality and Boldness represented a higher-level of personality organization. Correlations between personality factors and well-being were consistent with the definitions of the factors. There were no significant associations between personality and genotype. These results are consistent with the possibility that common marmoset personality structure varies as a function of rearing or housing variables that have not yet been investigated systematically.

## Introduction

Common marmosets (*Callithrix jacchus*) are small New-World monkeys that populate a wide range of habitats in South America [1, 2]. Because of their size, fast life history, and other physical and physiological characteristics, common marmosets are an increasingly popular animal model in biomedical research [3], although some [e.g., 4] have highlighted the shortcomings of marmoset models.

Common marmosets are also becoming popular subjects for research on cognition and personality. This trend has been driven partly by findings that common marmosets display behaviors and capabilities once believed to be exclusive to humans and great apes. Common marmosets, for example, exhibit high levels of spontaneous cooperative behavior [5, 6] and can discriminate between third parties that do and do not reciprocate [7]. These capabilities, and others, are believed to have evolved in common marmosets because they, like other callitrichids, but unlike other nonhuman primates, are cooperative breeders [see 8 for a review]. In species that are cooperative breeders, rather than disperse and mate, the adult siblings and offspring of mating pairs often stay within the family unit to help raise offspring, and so delay or forego reproduction [9].

Studies of common marmosets have revealed the presence of stable personality traits [10, 11], although one study found that these traits can be modified via social facilitation [12]. Studies have also found that personality traits in common marmosets are heritable and related to well-being [13], associated with the strength of laterality [14], and the binding potential of serotonin transporters in the brain [15]. Moreover, different methods, namely those based on behavioral observations and ratings, show evidence of convergent and discriminant validity in that they are correlated when both assess the same psychological construct, and uncorrelated when they do not, respectively [16, 17]. However, at least one study found that evidence for convergent and discriminant validity is not consistent across samples [16].

Studies of common marmosets have contributed to the attempt to reconstruct the evolutionary history of personality structure [18, 19]. Personality structure refers to the fact that statistical methods, including factor analysis and principal components analysis [20], but also others [e.g., 21], reveal that individual traits group into higher-order factors, which represent personality *domains*. In humans, for example, factor analysis has shown that traits such as ‘fearful’, ‘vulnerable’ and ‘anxious’ describe the Neuroticism domain while traits such as ‘active’, ‘social’, and ‘assertive’ describe the Extraversion domain [22, 23].

Like studies of other nonhuman primate taxa, including *Macaca* [24], *Pan* [25], *Saimiri* [26], *Sapajus* and *Cebus* [27-29], and other Callitrichids [30], studies of common marmosets [13, 17, 31, 32] have yielded findings consistent with the notion that a species’ socioecology influences its personality structure [see 33 for a discussion]. Despite differences in the origins and housing of subjects, and in how personality was measured, the five sets of data from these four studies revealed overlapping personality domains (see Table 1): all five revealed domains related to sociability [13, 17, 31, 32]; four revealed domains related to aggressiveness and competitive prowess [13, 17, 31, 32]; three revealed domains related to anxiety and vigilance [13, 17, 32]; two revealed domains related to exploratory tendencies [17, 31]; and two revealed domains related to self-control [17, 31]. In addition, two of these studies found a domain—Perceptual Sensitivity [17] and Patience [31]—that had not been found in other species, but which may also be related to self-control.

**Table 1.**
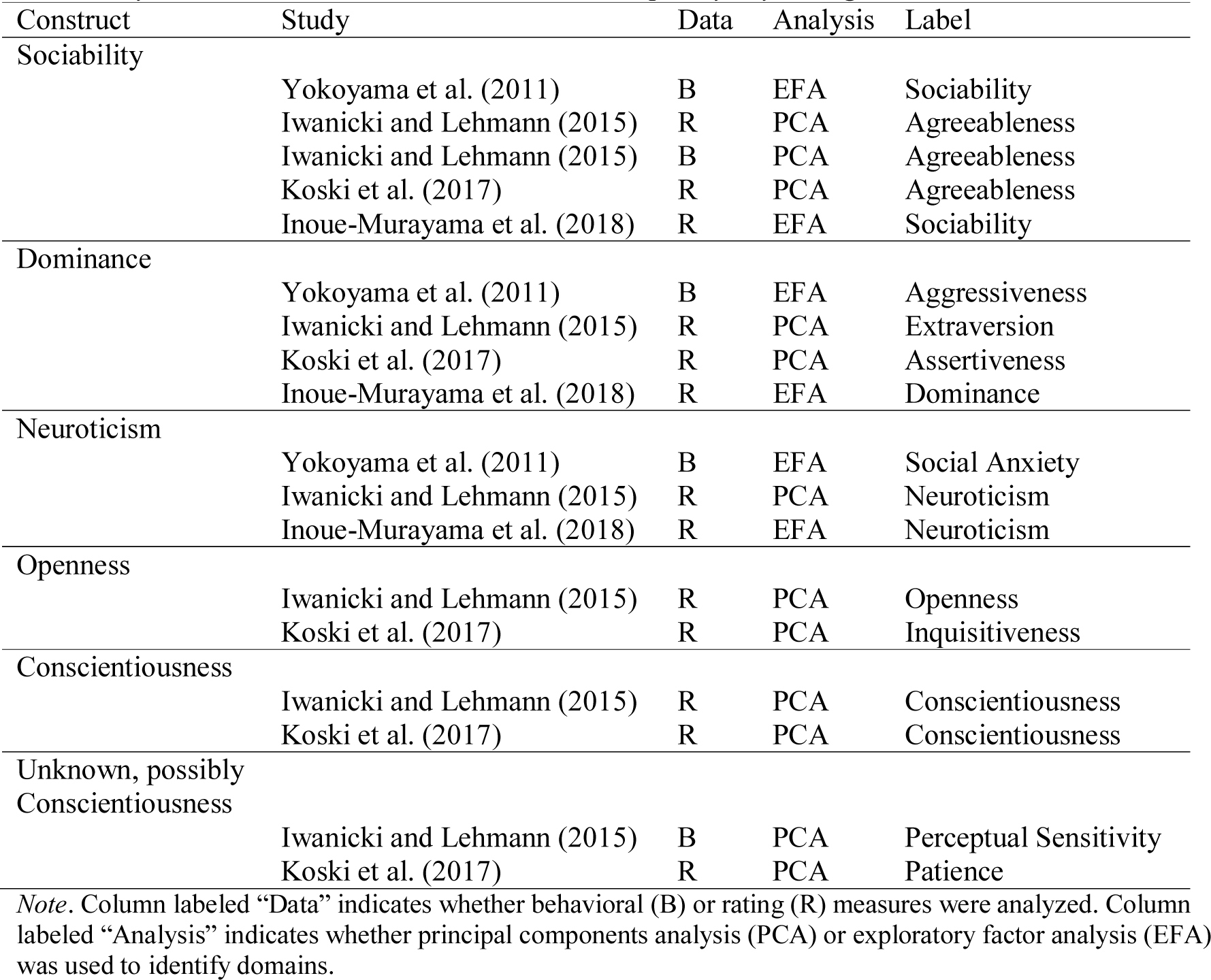
Personality Domains Found in Previous Studies Grouped by Psychological Construct

The most striking finding from these studies is that, although species exhibit individual differences in traits related to self-control [34, 35], in common marmosets these traits formed one or two broad domains. Among other primates, similar domains have only been found in humans *Homo sapiens* [e.g., 36], chimpanzees *Pan troglodytes* [37-42], bonobos *Pan paniscus* [25], and brown (or tufted) capuchin monkeys *Sapajus apella* [28], all of which are known for having larger brains. Consequently, findings of domains related to self-control in common marmosets led researchers to conclude that the cognitive and behavioral demands associated with cooperative breeding may have led to the evolution of these domains in what is a small-brained primate [31].

The results from the four studies of common marmosets, however, were not entirely consistent. Specifically, although they found evidence that common marmosets possess up to six or seven domains, they varied with respect to which subset of domains they found. This variation may be attributable to the fact that, asides perhaps from one study [31], these studies did not sample enough traits or individuals to capture all the ways in which individuals may differ in their personality.

To address whether this was the case, we followed up our earlier study of personality in 77 (68 male and 9 female) common marmosets housed at the Kobe, Japan campus of the Institute of Physical and Chemical Research (RIKEN) [13]. In that study, we did not find a Conscientiousness, Patience, or, for that matter, an Openness domain. One reason why we may not have found those domains is that several items related to these domains were removed because they had interrater reliability estimates that were less than zero [13]. This can happen if there is not enough between-subjects variance in a trait and/or a large amount of error variance [43].

A low level of between-subjects variance may come about when unmeasured influences, for example how animals are housed or bred, overwhelms, or makes it difficult to observe or perceive, individual differences in one or more traits. Recent findings that appear to rule out this possibility come from a study that found the same structure in laboratory- housed and wild common marmosets [44]. However, because the captive and wild groups lived in family groups, that study could not exclude the possibility that being raised in other kinds of groups might affect personality structure. A low level of between-subjects variance may also come about because the personalities of individuals conform to that of their group [12].

We attempted to rule out the possibility that low between-subjects variance was responsible for the inconsistency between our previous study that only found evidence for three domains—Dominance, Sociability, and Neuroticism—and the studies that found additional domains, including Conscientiousness. We therefore increased the between-subjects variance in the RIKEN sample by increasing the sample size by approximately two-thirds. Doing so also led to our nearly doubling the ratio of females to males.

In addition to trying to find these additional domains in the common marmosets housed at RIKEN, we examined associations between personality domains and subjective well-being. Previous studies in humans [45, 46] and in nonhuman primate species, including chimpanzees [40, 47, 48], orangutans *Pongo* spp. [49], rhesus macaques *Macaca mulatta* [50], brown capuchin monkeys [51], and common marmosets [13], have shown a consistent pattern of relationships between personality and measures of well-being or welfare. Specifically, personality domains associated with gregariousness, assertiveness, activity, and other traits associated with Extraversion [52] were related to higher subjective well-being and personality domains made up of traits associated with vigilance, fearfulness, anxiety, and other traits associated with Neuroticism [53] were associated with lower subjective well-being. By testing for associations between the personality domains and subjective well-being, then, we could assess the degree to which the personality domains we find are measures of distinct psychological constructs.

Finally, we tested whether a set of genetic polymorphisms were associated with personality. Our previous study of common marmosets found that lower Dominance and lower Neuroticism in common marmosets were both associated with the AA genotype of the μ-opioid receptor gene; lower Neuroticism was additionally associated with the short form of the arginine vasopressin receptor 1A gene [13]. For the present study we focused on single nucleotide polymorphisms (SNPs) of the serotonin receptor 1a gene. A study of chimpanzees identified a SNP (rs25209664: C743A) that caused a proline to glutamine substitution at the 248^th^ amino acid of the serotonin receptor 1a gene. This polymorphism was associated with aggression and sociability: chimpanzees who possessed two C alleles engaged in less social grooming and were rated as more anxious [54]. This study also found evidence for some interactions: males with the CC genotype displayed more often and, of chimpanzees with the AC genotype, mid-ranking individuals had lower proximity scores [54].

## Method

### Subjects

Subjects were 128 common marmosets (99 males, 29 females) that ranged in age from 1.6 to 15.1 (mean = 4.8, SD = 2.7). Subjects were recruited in three waves. The 77 subjects from the first wave had taken part in a similar study [13] and included 68 males and 9 females ranging in age from 1.5 to 15.1 years (mean=6.0, SD=2.6). Subjects from the second and third waves were born at RIKEN. The 24 subjects from the second wave included 17 males and 7 females ranging in age from 1.7 to 4.5 years (mean = 2.6, SD = 0.7) and the 27 subjects from the third wave included 14 males and 13 females ranging in age from 2.0 to 4.9 years (mean = 3.0, SD = 0.8).

### Animal Housing and Husbandry

Subjects were housed in the RIKEN Center for Biosystems Dynamics Research in Kobe, Japan. One hundred and twelve subjects were born at the center, six were supplied by CLEA Japan Inc. (Tokyo, Japan), and 10 were supplied by Japan Wild Animal Laboratory Limited (Amami, Japan). Subjects sourced from other facilities had lived in the center for at least three years prior to this study.

At RIKEN, subjects were housed in breeding rooms that had a 12 h light-dark cycle (light: 08:00–20:00). Enclosures (1630 × 760 × 831 mm for families, 660 × 650 × 600 or 660 × 450 × 600 mm for pairs or individuals) had wooden perches, a plastic cube-shaped shelter, a food tray, and a water dispenser. There were around twenty cages in each breeding room and so even if animals were individually housed, they were exposed to visual, auditory, and olfactory stimulation from conspecifics. The temperature and humidity in the breeding room were maintained at approximately 28°C and 50%, respectively. In the morning and afternoon, subjects received solid food (CMS-1, CLEA Japan, Inc., Tokyo, Japan) mixed with an appropriate amount of water to soften it, powdered milk formula, honey, gluconic acid, calcium, vitamin C, and lactobacillus probiotic. Food in the afternoon was softened into a paste by soaking it in water and then stirring it. Once a week subjects’ diets were supplemented with chopped and boiled eggs or bananas.

### Animal Rearing

Animals reared by their parents and/or their family members, including one to five older brothers or sisters, were reared in family cages. At around 14 days after birth, when they were still infants, these subjects were fed a food paste in the afternoon. When these individuals were between 6 and 15 months old, to ensure that they were provided with the required amount of space, they were transferred from their family cage to a home cage (0.21 to 0.43m^2^ floor space per animal). Animals living in these home cages were same-sex, mixed-age peers. Individuals that were to be used in brain imaging [55] or in behavioral studies, and individuals that did not get along with their partners, were single housed.

Animals that were not reared by their parents, for example, in the event of a triplet birth or parental neglect, were hand-reared in climate-controlled rearing cages by human caregivers. This procedure has been described elsewhere [13]. In short, these animals were housed in a thermal insulation box and a towel roll from one day to 21 days after birth. Then, from 21 days after birth to weaning, these animals were housed in a wire-mesh box sized 390 × 230 × 300 mm furnished with a hammock, perches, towel roll, feeding dish, and water bottle. These animals were breastfed on the day of their birth and then bottle-fed until weaning. A food paste was introduced at around 28 days and then animals were weaned fully 50 to 70 days after birth. After weaning, these animals were housed in a home cage with peers or individually in the breeding room.

Of the 77 subjects from the first wave, 30, including 23 parent-reared and seven hand-reared subjects, were housed in a family group (n = 13) or with same-sex peers (n = 17). The remaining 47 subjects, including 33 that were parent-reared, 13 that were hand-reared, and one with an unknown rearing history, were single-housed. Of the 24 subjects from the second wave, 22, including 18 that were parent-reared, 3 that were hand-reared, and one with an unknown rearing history, were housed in a family group (n = 7), with an opposite sex marmoset for breeding (n = 2), or with same-sex peers (n = 13). The remaining two subjects from the second wave were parent-reared and single-housed. Of the 27 subjects from the third wave, 25 parent-reared subjects and one hand-reared subject were housed in a family group (n = 1), with an opposite-sex marmoset for breeding (n = 4), with same-sex peers (n = 19), or single-housed (n = 2). The remaining subject was parent-reared and single-housed.

### Ratings

#### Questionnaires

For this study, we used versions of the personality and subjective well-being questionnaires that had been translated into Japanese using a back-translation procedure. A study of chimpanzees revealed that the translation did not affect the psychometric properties of these questionnaires [40].

**Personality.** To assess personality, we used the Hominoid Personality Questionnaire (HPQ).^1^ Each of the HPQ’s 54 items consists of a trait adjective paired with one to three sentences that set the adjective in the context of primate behavior. For example, “**FEARFUL**” (boldface and capitals in the original) is paired with the descriptor sentence “Subject reacts excessively to real or imagined threats by displaying behaviors such as screaming, grimacing, running away or other signs of anxiety or distress.” The HPQ’s instructions ask raters to **a)** judge the standing of each animal on each trait based on the animal’s behavior and interactions with others, and the rater’s own judgement of what constitutes average behavior for this species, **b)** assign a rating of 1 (“Displays either total absence or negligible amounts of the trait.”) to 7 (“Displays extremely large amounts of the trait.”) to each item, and **c)** not discuss their ratings with their fellow raters.

A description of the HPQ’s development can be found elsewhere [56]. Briefly, the HPQ grew out the 48-item Orangutan Personality Questionnaire [49], which grew out of the 43-item Chimpanzee Personality Questionnaire [39]. Forty-one of the HPQ’s 54 items were sampled from Goldberg’s [57] trait terms of the five major domains of human personality [39]. The remaining 13 items were adapted from items [58] or facets [59] from other human personality inventories, or were created for by the authors of these instruments [39, 40, 49].

**Subjective Well-Being.** Ratings were made on a four-item scale that was based on a questionnaire used to measure subjective well-being in captive chimpanzees [47]. Each item was devised to assess a different concept of subjective well-being that had been described in the human literature [47, 60-64]. The first item (moods) concerned the extent to which an individual experienced positive versus negative affect. The second item (social) concerned whether the individual experienced pleasure from social interactions. The third item (goals) concerned whether the individual was able to achieve its goals, bearing in mind that different individuals may have different, personal goals. The fourth item (be marmoset) asked raters how “happy” they would be if they were that marmoset for a week and was thus meant to measure global satisfaction. The subjective well-being scale’s instructions asked raters to assign a rating of 1 (“Displays either total absence or negligible amounts of the trait or state.”) to 7 (“Displays extremely large amounts of the trait.”) to each item. The instructions also request that raters do not discuss their ratings.

#### Raters and Ratings

We asked three keepers (two men and one woman) who completed the questionnaires in the first wave of data collection [see 13 for details] to complete the questionnaires for the second and third wave of data collection. The keepers did not know the results of the previous study or the purpose of collecting the data. The keepers had known the subjects they rated for 1.1 to 9.8 years (mean = 3.7 years, SD = 2.2). Two keepers (one man and one woman) rated all 128 subjects and the third rated 81 subjects. This resulted in a total of 337 ratings or an average of 2.63 ratings per subject. There were no missing rating data.

### Genotyping

A buccal swab was taken from each subject and kept in a 90% ethanol solution until DNA extraction. DNA was extracted by DNeasy Blood and Tissue kit (Qiagen, CA, USA). PCR amplification was conducted in a 10 μl (the total volume) reaction mixture containing 10ng of DNA template, 0.4 µM of each primer (forward: 5’-tggattcccttcctccgaaa-3’, reverse: 5’-aggtgttgattccctagggt-3’), 0.5U of LA Taq DNA polymerase, 400 µM of dNTPs, and GC buffer I (TaKaRa, Shiga, Japan). After denaturing DNA samples at 95°C for 1 min, we set up 40 cycles of 95°C for 30 seconds, 60°C for 30 seconds, 74°C for 1 minute, and a final extension at 74°C for 10 minutes. A total of 1,473 base pair fragments including whole single exon region were amplified. We then sequenced the polymerase chain reaction products, both forwards and backwards, using 3130xl Genetic Analyzer (Applied Biosystems, CA, USA). The internal primer 5’-tcatgctggttctctatggg-3’ was also used for sequencing. Primers were designed based on the NCBI Reference Sequence NC_013897. In the end, we identified three novel SNPs (G840C, G841A, and T901A) in the third intracellular region of the receptor (see Figures 1 and 2). G840C was a synonymous SNP coding alanine at the 280^th^ amino acid sequence, G841A was a nonsynonymous SNP that caused a methionine substitution at the 281^st^ amino acid sequence, and T901A was a nonsynonymous SNP that caused a serine to threonine substitution at the 301^st^ amino acid sequence.

**Figure 1.**
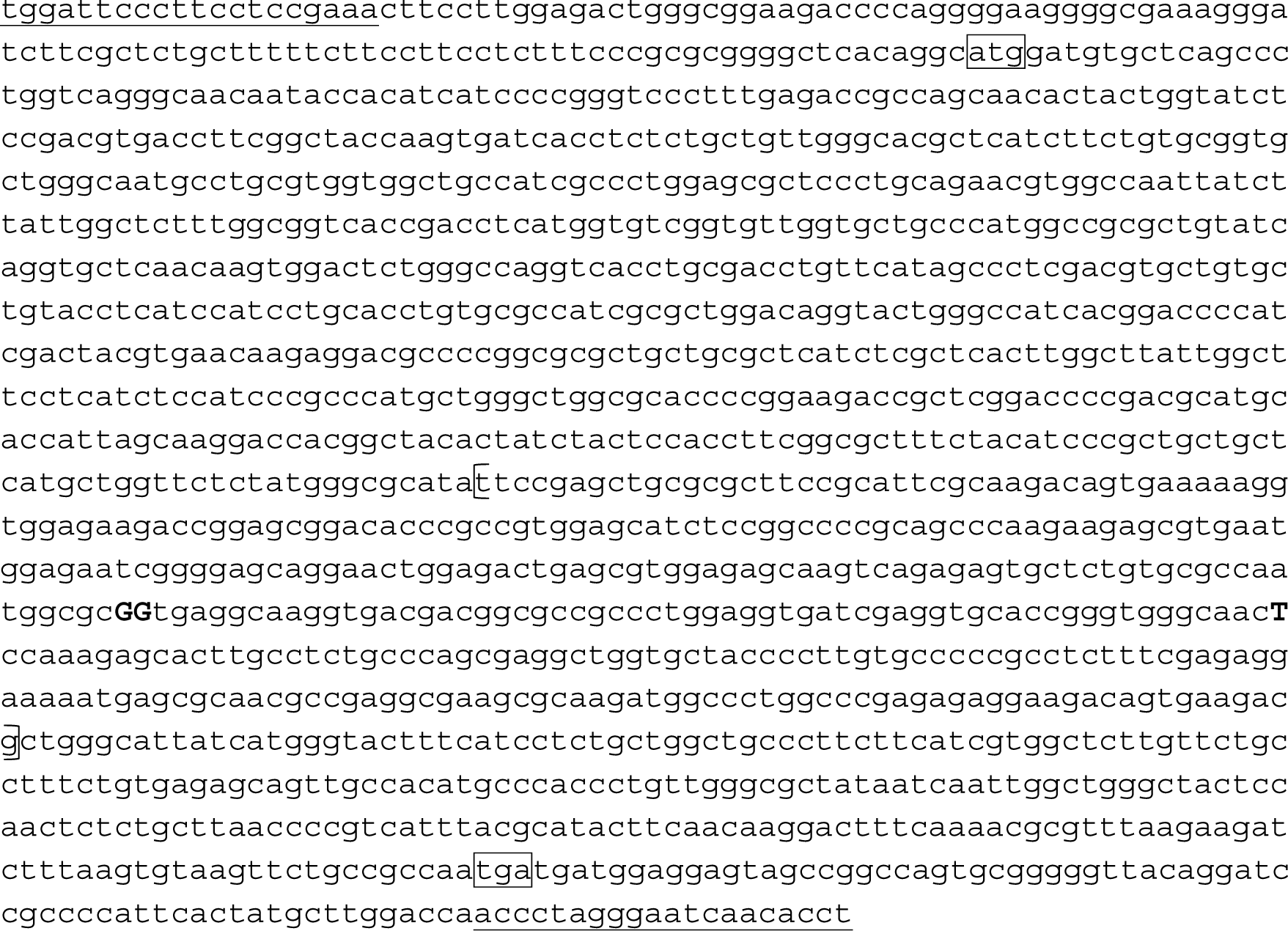
The sequence is based on NC_013897. Primer sequences are underlined. Start and stop codon sequences are in boxes. The third intracellular region is enclosed in the parentheses. Nucleotide substitutions are shown in capital and bold letters; G840C (A280A), G841A (V281M), T901A (S301T).

**Figure 2.**
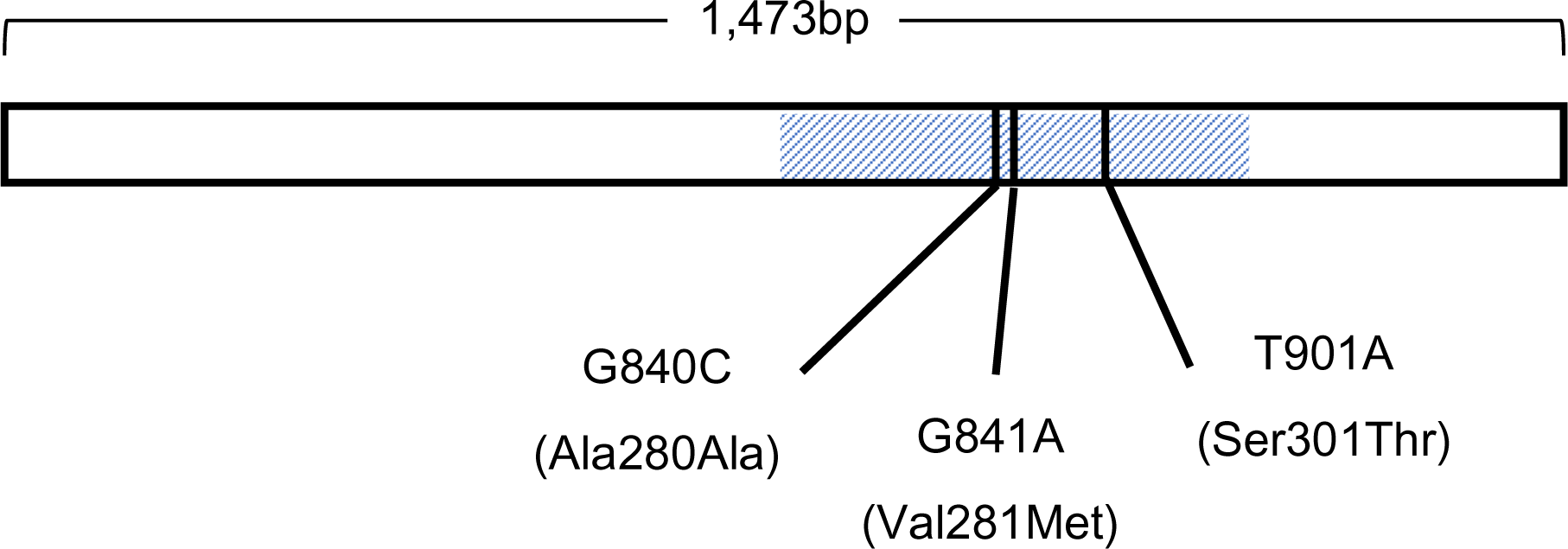
Locations of the single nucleotide polymorphisms on the single exon of the common marmoset serotonin receptor 1a gene *Note.* The shaded region corresponds to the third intracellular region.

### Analyses

We conducted the analyses using version 3.6.3 of R [65]. We used functions from version 1.9.12 of the psych package [66], version 1.0.7 of the EFA.MRFA package [67], and some custom functions.

#### Item Interrater Reliabilities

We used a custom function to compute the interrater reliabilities of the ratings of individual items on the HPQ and subjective well-being questionnaire. This function computed two intraclass correlations (*ICC*s) described by Shrout and Fleiss (43). One, *ICC*(3,1), indicates the reliability of individual ratings, that is, it is an estimate of the reliability of the rating from a single rater. The other, *ICC*(3,*k*), indicates the reliability of the mean rating coming from *k* raters, which was equal to 2.63 in the present study. We excluded items that had reliabilities that were less than or equal to zero [see 56 for a discussion].

#### Exploratory Factor Analyses

**Personality.** We conducted these analyses on the aggregate (mean) of personality ratings for the 128 subjects. Simulation studies [68-70] have shown that the number of subjects required for satisfactory recovery of factors is a function of item communalities, item loadings, and the item:factor ratio, and that the subject:item ratio is irrelevant. Previous rating-based studies of common marmoset personality [13, 17, 31] found that 72% to 97% of questionnaire items were reliable, a total of three to five factors, median salient loadings of around .6 or .7, and communalities that were between wide and high [see ref 69 for definitions of these types of communalities]. As such, we have a large enough sample size to conduct factor analyses on these data [69].

Before extracting factors using the maximum likelihood procedure, we determined how many factors to extract. To do so, we used the fa.parallel function from the psych package to generate a scree plot, which we inspected, and to conduct a parallel analysis [71] in which we compared eigenvalues from a principal components analysis of our data to the distribution of 1000 eigenvalues generated from principal components analysis of resampled and randomly generated data. We used principal components analysis for our parallel analysis because a recent study showed that the number of dimensions identified in this manner is more accurate [72]. In addition, we used the VSS function from the psych package to determine, for one to eight factor solutions, which had the lowest Bayesian Information Criterion [BIC; 73], and the hullEFA function from the EFA.MRFA package to determine the number of factors via the Hull method [74], which is known to perform well with personality data [75]. Finally, we inspected the factors to ensure that they were interpretable.

After we extracted factors using the fa function from the psych package, we applied an oblique (promax) and an orthogonal (varimax) rotation. If the promax rotation yielded factors that were strongly correlated and/or a different structure, we retained and interpreted those factors. We otherwise retained and interpreted the varimax-rotated factors.

In interpreting factors, we specified that salient loadings were those equal to or greater than |0.4|. We labeled factors based on our interpretations of what they were and attempted to find suitable labels from previous studies of common marmosets, and, if none were available, studies of nonhuman primates and humans. If we could not find a label from these sources, we devised our own.

In addition to conducting this first-order factor analysis, we found evidence (results to be discussed) suggesting that there may be second-order personality factors underlying these data. We thus conducted an exploratory factor analyses of the inerfactor correlation (Phi) matrix obtained from the promax-rotated factors. For the same reason, we conducted an additional item-level factor analysis as a robustness check.

**Subjective Well-Being.** We conducted a maximum likelihood factor analysis using the fa function on aggregated (mean) ratings for all 128 subjects. Previous work in 77 of these subjects revealed a single factor [13].

#### Unit-Weighted Factor Scores

For the remaining analyses, we used a custom function to compute unit-weighted scores [20, 76] for the personality and subjective well-being data. This involved, for each item, finding the largest salient factor loading. If that loading was positive, we assigned it a weight of +1. If that loading was negative, we assigned it a weight of -1. In all other cases, we assigned a weight of zero. After We then summed the weighted item ratings.

#### Factor Reliabilities

For the first- and second-order personality factors, and for subjective well-being, we used the same custom function to compute *ICC*(3,1) and *ICC*(3,*k*) for the items to compute these *ICC*s. As with the item-level analyses, *k* was equal to 2.63. In addition, we used the alpha function from the psych package to compute Cronbach’s alpha (α), a measure of the internal consistency reliability of a scale, and the omega function to compute McDonald’s omega hierarchical (ω_h_), a measure of the degree to which a general factor saturates a scale’s items.

#### Personality Factor Comparisons

To compare the first- and second-order factors to factors found in previous studies of common marmoset personality [13, 17, 31], we first generated unit-weighted factor scores based on the personality structures described in these other studies. Because there was not a total overlap of questionnaire items across these studies, we sometimes substituted items that were similar in their meaning or in the constructs that they purportedly assessed. Details about how these scores were created can be found in Table S1. After computing these unit-weighted scores, we obtained correlations between the scores based on the factor loadings from the present study and the scores based on component and factor loadings from previous studies. We compared the absolute magnitudes of these correlations and highlighted the highest correlation or, in the case where the confidence intervals of two or more correlations overlapped, highest correlations.

#### Personality-Subjective Well-Being Associations

We used Pearson correlation coefficients to examine associations between the first- and second-order personality factors and, both, the subjective well-being items and the sum of these items. We used Holm’s method [77] to adjust for familywise error rates.

#### Genetic Associations

To examine the genotype-personality associations, for the first- and second-order personality factors, we fit linear models using the lm function. For these analyses, we standardized the personality factor scores (mean = 0, SD = 1). The variables in the models included sex (male = 1, female = 0), age in years, and a categorical variable that indicated genotype. Because there were problems with genotyping four subjects, these individuals were excluded from the analyses. In addition, the G840C genotypes for two subjects and T901A genotype for one subject were unclear, and so these individuals were not included in tests of associations between personality and the G840C and T901A genotype, respectively. Finally, only one subject had the AA version of G841A and only nine subjects had the GA version of this genotype. We therefore did not examine associations between these genotypes and personality.

Although subjects were related, we did not test for the effect of genotypes within the context of an animal model [cf. 13]. Moreover, because we conducted multiple, sometimes non-independent, tests, we used the Bonferroni correction to adjust for familywise error rates.

### Ethics

This study complied with the current laws of Japan, including the Act on Welfare and Management of Animals. All experimental and husbandry procedures were performed in accordance with RIKEN’s Guidelines for Conducting Animal Experiments, and in accordance with the ARRIVE (Animal Research: Reporting of In Vivo Experiments) guidelines. All procedures were approved by the Animal Care and Use Committee of the Kobe Institute of RIKEN (MA2009-10-16).

## Results

### Interrater Reliabilities of Items

#### Personality

The interrater reliabilities of the 54 HPQ items are presented in Table 2. The reliabilities of individual ratings and of mean ratings for the items ‘anxious’, ‘persistent’, ‘quitting’, and ‘unperceptive’ were negative, and so we excluded these items from further analyses. Although the reliability of mean ratings for the items ‘innovative’ and ‘decisive’ were equal to 0.01, the reliabilities of individual ratings for these items were less than 0.01, and so we decided to exclude those items, too.

**Table 2.** Interrater Reliabilities of Items from the Hominoid Personality Questionnaire

| Item | *ICC(3,1)* | *ICC(3,k)* |
| --- | --- | --- |
| Sociable | 0.45 | 0.68 |
| Sympathetic | 0.43 | 0.66 |
| Solitary | 0.43 | 0.67 |
| Protective | 0.42 | 0.66 |
| Helpful | 0.38 | 0.62 |
| Friendly | 0.36 | 0.60 |
| Aggressive | 0.34 | 0.57 |
| Gentle | 0.34 | 0.58 |
| Irritable | 0.33 | 0.57 |
| Affectionate | 0.32 | 0.55 |
| Stingy/greedy | 0.29 | 0.51 |
| Dominant | 0.28 | 0.50 |
| Individualistic | 0.28 | 0.51 |
| Submissive | 0.28 | 0.51 |
| Imitative | 0.28 | 0.51 |
| Independent | 0.27 | 0.50 |
| Excitable | 0.26 | 0.48 |
| Conventional | 0.26 | 0.48 |
| Bullying | 0.23 | 0.44 |
| Impulsive | 0.23 | 0.44 |
| Dependent/follower | 0.23 | 0.44 |
| Defiant | 0.23 | 0.45 |
| Stable | 0.21 | 0.41 |
| Jealous | 0.21 | 0.41 |
| Cautious | 0.19 | 0.39 |
| Active | 0.19 | 0.38 |
| Fearful | 0.18 | 0.37 |
| Autistic | 0.18 | 0.36 |
| Thoughtless | 0.17 | 0.36 |
| Erratic | 0.17 | 0.35 |
| Playful | 0.16 | 0.33 |
| Intelligent | 0.16 | 0.33 |
| Reckless | 0.15 | 0.32 |
| Curious | 0.14 | 0.30 |
| Sensitive | 0.14 | 0.30 |
| Inquisitive | 0.13 | 0.28 |
| Cool | 0.13 | 0.29 |
| Predictable | 0.13 | 0.28 |
| Lazy | 0.11 | 0.25 |
| Timid | 0.10 | 0.22 |
| Clumsy | 0.09 | 0.21 |
| Disorganized | 0.09 | 0.20 |
| Vulnerable | 0.07 | 0.16 |
| Unemotional | 0.06 | 0.15 |
| Distractible | 0.05 | 0.13 |
| Manipulative | 0.05 | 0.12 |
| Depressed | 0.04 | 0.09 |
| Inventive | 0.02 | 0.06 |
| Innovative | 0.00 | 0.01 |
| Decisive | 0.00 | 0.01 |
| Anxious | -0.03 | -0.08 |
| Persistent | -0.06 | -0.19 |
| Quitting | -0.08 | -0.24 |
| Unperceptive | -0.12 | -0.39 |

Of the remaining 48 items, the interrater reliabilities of individual ratings ranged from 0.02 (‘inventive’) to 0.45 (‘sociable’). The mean and standard deviation for these estimates were 0.21 and 0.11, respectively. The interrater reliabilities of mean ratings for the remaining items ranged from 0.06 (‘inventive’) to 0.68 (‘sociable’). The mean and standard deviation for these estimates were 0.40 and 0.16, respectively.

#### Subjective Well-Being

The interrater reliabilities of individual ratings were 0.16, 0.10, 0.15, and 0.14 for the moods, social, goals, and global well-being items, respectively. The corresponding reliabilities of mean ratings were 0.33, 0.23, 0.32, and 0.30.

### Maximum Likelihood Exploratory Factor Analyses

#### Personality

**First-Order Analysis.** The scree plot (see Figure S1) and parallel analysis indicated that there were five factors. The Hull method (see Figure S2) also indicated that there were five factors, and BIC achieved a minimum with five factors. We therefore extracted five factors.

A promax rotation of the five-factor solution yielded two interfactor correlations that were large (*r*s ≥ |0.5|) and two that were medium-sized (*r*s ≥ |0.3|). The mean and standard deviation of the absolute interfactor correlations were 0.26 and 0.20, respectively. Comparison of the varimax- and promax-rotated factors revealed that the congruence coefficients for two factors fell below 0.95 (see Table S2) and an inspection of the loadings indicated that the promax-rotated factors differed some from their varimax-rotated counterparts. Given these results, we interpreted the promax-rotated factors (see Table 3), which explained 63% of the variance (the varimax-rotated factors are presented in Table S3).

**Table 3.** Pattern Matrix from the First-Order Factor Analysis of the Hominoid Personality Questionnaire

| Item | Factor Loadings | | | | | $h^2$ |
| --- | --- | --- | --- | --- | --- | --- |
|  | Soc | Dom | Imp | Opn | Neg |  |
| Helpful | **0.89** | 0.10 | -0.05 | 0.09 | -0.08 | 0.79 |
| Sympathetic | **0.83** | -0.01 | -0.07 | 0.07 | 0.08 | 0.76 |
| Protective | **0.80** | 0.01 | 0.02 | 0.02 | -0.15 | 0.66 |
| Individualistic | **-0.78** | 0.09 | -0.05 | 0.13 | 0.25 | 0.71 |
| Dependent/follower | **0.77** | 0.06 | -0.05 | 0.23 | 0.39 | 0.69 |
| Independent | **-0.73** | 0.27 | -0.31 | 0.10 | 0.08 | 0.62 |
| Imitative | **0.72** | 0.04 | 0.07 | 0.20 | 0.18 | 0.49 |
| Solitary | **-0.71** | 0.05 | -0.04 | -0.17 | 0.34 | 0.74 |
| Affectionate | **0.64** | -0.09 | -0.16 | 0.21 | 0.12 | 0.65 |
| Sensitive | **0.64** | -0.08 | -0.17 | -0.02 | 0.04 | 0.61 |
| Sociable | **0.62** | -0.27 | -0.16 | 0.16 | -0.15 | 0.85 |
| Conventional | **0.60** | 0.11 | -0.33 | -0.13 | 0.19 | 0.62 |
| Intelligent | **0.59** | 0.26 | -0.18 | 0.00 | -0.19 | 0.39 |
| Gentle | **0.50** | -0.35 | -0.24 | 0.15 | 0.09 | 0.83 |
| Reckless | **-0.50** | -0.16 | 0.39 | 0.38 | -0.03 | 0.62 |
| Friendly | **0.49** | **-0.46** | -0.15 | 0.13 | 0.02 | 0.86 |
| Jealous | 0.02 | **0.92** | -0.04 | 0.21 | 0.15 | 0.83 |
| Stingy/greedy | -0.08 | **0.87** | -0.06 | 0.28 | 0.15 | 0.86 |
| Bullying | -0.04 | **0.85** | 0.00 | 0.11 | 0.07 | 0.77 |
| Dominant | -0.09 | **0.79** | 0.06 | 0.01 | -0.07 | 0.80 |
| Manipulative | 0.21 | **0.72** | -0.10 | 0.04 | -0.33 | 0.57 |
| Aggressive | -0.12 | **0.69** | 0.15 | -0.05 | -0.17 | 0.80 |
| Defiant | -0.09 | **0.65** | 0.20 | -0.01 | -0.20 | 0.78 |
| Irritable | 0.00 | **0.50** | **0.48** | -0.18 | -0.11 | 0.74 |
| Excitable | -0.02 | 0.16 | **0.78** | -0.08 | -0.06 | 0.78 |
| Unemotional | -0.10 | 0.15 | **-0.77** | 0.00 | 0.26 | 0.52 |
| Impulsive | -0.10 | 0.06 | **0.75** | 0.13 | 0.13 | 0.76 |
| Cool | 0.18 | -0.05 | **-0.66** | -0.06 | 0.01 | 0.65 |
| Fearful | 0.23 | -0.10 | **0.62** | **-0.42** | 0.28 | 0.51 |
| Distractible | -0.11 | -0.05 | **0.53** | 0.23 | 0.10 | 0.40 |
| Disorganized | -0.10 | 0.15 | **0.52** | 0.18 | 0.12 | 0.51 |
| Stable | 0.21 | -0.19 | **-0.46** | 0.07 | -0.38 | 0.64 |
| Thoughtless | -0.22 | -0.03 | **0.41** | 0.38 | 0.02 | 0.45 |
| Predictable | 0.12 | -0.08 | **-0.40** | 0.04 | 0.02 | 0.27 |
| Erratic | -0.20 | 0.32 | 0.35 | -0.06 | 0.20 | 0.54 |
| Curious | 0.14 | 0.13 | 0.08 | **0.73** | 0.02 | 0.61 |
| Inquisitive | 0.17 | 0.11 | 0.06 | **0.72** | 0.09 | 0.55 |
| Inventive | 0.27 | 0.18 | -0.05 | **0.68** | 0.11 | 0.51 |
| Playful | 0.23 | -0.12 | 0.32 | **0.65** | -0.04 | 0.60 |
| Cautious | **0.47** | 0.08 | 0.24 | **-0.59** | 0.21 | 0.53 |
| Active | 0.28 | 0.18 | **0.41** | **0.52** | -0.13 | 0.69 |
| Autistic | 0.03 | -0.13 | 0.09 | 0.19 | **0.68** | 0.45 |
| Timid | 0.14 | 0.00 | 0.34 | -0.18 | **0.65** | 0.58 |
| Depressed | -0.17 | 0.07 | -0.27 | -0.04 | **0.64** | 0.51 |
| Clumsy | -0.04 | 0.17 | -0.05 | 0.00 | **0.58** | 0.33 |
| Vulnerable | -0.02 | -0.19 | -0.02 | 0.04 | **0.57** | 0.39 |
| Lazy | -0.23 | -0.05 | **-0.45** | -0.17 | **0.50** | 0.59 |
| Submissive | 0.33 | -0.20 | -0.18 | -0.01 | **0.48** | 0.61 |
| Proportion of variance | 0.20 | 0.14 | 0.13 | 0.08 | 0.08 |  |

| Factor | Factor Correlations |  |  |  |  |
| --- | --- | --- | --- | --- | --- |
|  | Soc | Dom | Imp | Opn | Neg |
| Soc | 1.00 |  |  |  |  |
| Dom | -0.53 | 1.00 |  |  |  |
| Imp | -0.45 | 0.57 | 1.00 |  |  |
| Opn | 0.06 | 0.15 | 0.18 | 1.00 |  |
| Neg | -0.06 | -0.19 | -0.04 | -0.38 | 1.00 |

*Note.* $N = 128$ . Soc = Sociability, Dom = Dominance, Imp = Impulsiveness, Opn = Openness, Neg = Negative Affect, $h^2$ = communalities. Factors extracted using a maximum likelihood estimation and rotated using the promax procedure. Factor loadings greater than or equal to $|0.4|$ are in bold.

The first factor loaded on items related to high Extraversion and high Agreeableness in humans [e.g., 57]. This factor resembled the Sociability factor in one study [13] and the Agreeableness factor from two other studies [17, 31] of common marmosets. To be consistent with our prior study [13], we labeled this factor Sociability.

The second factor loaded on items related to high Extraversion and low Agreeableness in humans [e.g., 57]. Compared to other studies of common marmosets, it is best described as a narrow version of factors labeled Dominance [13], Extraversion [17], and Assertiveness [31]. To be consistent with our prior study [13], we labeled this factor Dominance.

The third factor had positive loadings on items related to high Neuroticism and low Conscientiousness in humans, and also negative loadings on items related to low Neuroticism and high Conscientiousness [e.g., 57]. In a previous study of a subsample of these animals [13], the Dominance and Neuroticism factor loaded on some of these items. Compared to other studies of common marmosets, it resembled most closely the factors labeled Conscientiousness and Patience in one study [31] and the Conscientiousness factor in another [17]. Humans that are high in Neuroticism and low in Conscientiousness are described as having an undercontrolled style of impulse control [78]. We thus labeled this factor Impulsiveness.

With the exception of a negative loading on the item cautious, the fourth factor loaded primarily on items related to high Openness in humans [e.g., 57]. Previous studies of common marmosets have labeled factor such as these Openness [17] and Inquisitiveness [31]. We therefore labeled this factor Openness.

The fifth factor loaded on items related to low Extraversion and high Neuroticism in humans [e.g., 57]. In the previous study of a subset of these animals [13], Neuroticism had a positive loading on many of these items. In all three previous studies of this species, factors such as Dominance, Assertiveness, and Conscientiousness had negative loadings on these items [13, 17, 31]. Given that this factor combined aspects of high Neuroticism and low degrees of Assertiveness or social prowess, we labeled it Negative Affect.

**Second-Order Analysis.** Because there were several non-negligible correlations between the just-described factors, we factor analyzed the factor intercorrelation (Phi) matrix. Inspection of the scree plot (see Figure S3) and parallel analysis indicated that there were two factors; BIC was lowest for the two-factor solution.^2^ We also tried to extract a single ‘general’ factor, but this solution exhibited poor fit (root mean square of the residuals = 0.14), and the factor did not have a salient loading on Openness or on Negative Affect. A promax rotation of the two-factor solution indicated that they were close to being orthogonal, and the loadings of the varimax-rotated factors were nearly identical to those of the promax-rotated factors (see Table 4). We therefore interpreted the varimax-rotated factors, which accounted for 49% of the variance. After reflecting (multiplying loadings by -1) the first factor, it had a positive loading on Sociability and a negative loading on both Dominance and Impulsiveness. We thus labeled this factor Pro-sociality. The second factor had a positive loading on Openness and a negative loading on Negative Affect. We thus labeled this factor Boldness.

**Table 4.** Pattern Matrix from the Second-Order Factor Analysis of Personality Factors

| First-Order Factor | Promax Rotation | | Varimax Rotation | | $h^2$ |
| --- | --- | --- | --- | --- | --- |
|  | Pro-sociality | Boldness | Pro-sociality | Boldness |  |
| Dominance | **-0.79** | 0.15 | **-0.80** | 0.21 | 0.68 |
| Sociability | **0.72** | 0.21 | **0.70** | 0.15 | 0.51 |
| Impulsiveness | **-0.67** | 0.08 | **-0.68** | 0.13 | 0.48 |
| Negative affect | -0.01 | **0.62** | 0.04 | **-0.61** | 0.38 |
| Openness | -0.01 | **-0.61** | -0.06 | **0.61** | 0.38 |
| Proportion of variance | 0.32 | 0.17 | 0.32 | 0.17 |  |

*Note.* $N = 128$ . $h^2$ = communalities. Factors extracted from the factor correlation matrix from Table 2 using a maximum likelihood estimation and rotated using the promax and varimax procedures. The Pro-Social factor for both structures were reflected, that is, the loadings were multiplied by -1. The promax-rotated factors correlated -0.15. Factor loadings greater than or equal to $|0.4|$ are in bold.

**Robustness Checks.** Previous studies of common marmosets did not find evidence higher-order factors, that is, interfactor correlations tended to be modest [13, 17, 31]. We thus felt it would be worthwhile to follow-up these results.

The first of our robustness checks tested whether the higher-order factors reflected the structuring of data collection. Specifically, because different subjects were rated in each wave, this may have led raters to rate the subjects belonging to each wave as resembling one another more than they did subjects in other waves. To test this, we residualized the 48 reliable items on a categorical variable that represented whether the subject was rated in the first, second, or third wave. We then factor analyzed the residualized item scores.

Inspection of the scree plot (see Figure S4) suggested that there were four or five factors and parallel analysis indicated that there were four factors. The Hull test indicated that there were two factors (see Figure S5). The BIC was lowest for a five-factor solution. We therefore examined promax-rotations after extracting two, four, and then five factors.

For the two-factor solution (see Table S4), the first factor loaded predominantly on items onto which the factors Sociability, Dominance, and Impulsiveness had loaded. It therefore resembled the higher-order factor Pro-sociality. The second factor loaded predominantly on items onto which the factors Openness and Negative Affect had loaded. It therefore resembled the higher-order factor Boldness. For the four-factor solution (see Table S5), the first, third, and fourth factors resembled Sociability, Negative Affect, and Openness. The second factor loaded predominantly on items related to high Dominance and high Impulsiveness. The five-factor solution yielded five factors that resembled the five factors that had been found earlier (see Table S6).

The similarity, as indicated by Tucker’s congruence coefficients, between the five factors obtained before and after item scores were residualized were equal to or greater than 0.98, suggesting that these were the same factors (see Table S7). Factor analysis of the residualized item scores, then, revealed either the same structure (the five-factor solution) or structures in which there were stronger associations between factors (the two- and four-factor solutions). These results are not consistent with the possibility that the higher-order factors reflect the fact that we collected these data in three stages.

The second of our robustness checks was to test whether the second-order factors are general evaluative factors used by raters [cf. 79]. To do so, we factor analyzed ratings from each of the three raters separately. We also factor analyzed a weighted correlation matrix from which removed possible effects of raters:

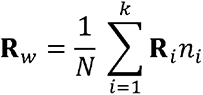

where **R***_w_*, the weighted correlation matrix, is the sum of the products of the correlation matrices of each of *k* = 3 raters, **R**_i_, and the subjects, *n*_i_ rated by that individual rater, divided by the total number of subjects, *N*.

For ratings from one keeper who rated all the subjects, the scree plot (see Figure S6), parallel analysis, BIC, and Hull method (see Figure S7) indicated that there were five factors. We thus extracted five factors and subjected them to a promax rotation (see Table S8). These factors resembled those obtained in the initial factor analysis and the interfactor correlations were similar in magnitude. We then conducted a second-order factor analysis in which we forced a two-factor solution. Although one second-order factor just missed our criterion for a salient loading on a first-order factor, these factors resembled Pro-sociality (reversed) and Boldness (see Table S9).

For ratings from the second keeper who rated all the subjects, the scree plot (see Figure S8) indicated that there were five factors and parallel analysis, the BIC, and the Hull method (see Figure S9) indicated that there were four factors. We thus extracted four factors and subjected the m to a promax rotation (see Table S10). The first factor appeared to be a Dominance versus Agreeableness, the third factor was Gregariousness (a narrow facet of Extraversion), and the last two factors were difficult to interpret. The interfactor correlations were not consistent with there being a second-order factor.

For ratings from the keeper who rated 81 subjects, the scree plot (see Figure S10) and parallel analysis, the BIC, and Hull method (see Figure S11) indicated that there were three factors. We thus extracted three factors and subjected them to a promax rotation (see Table S11). These factors included Agreeableness versus Dominance, Extraversion, and Negative Affect/Impulsiveness, respectively.

For the weighted correlation matrix, the scree plot indicated that there were five factors (see Figure S12) and both the parallel analysis, and the BIC indicated that there were three factors. We thus extracted three factors and subjected them to a promax rotation (see Table S12). The first and third factors loaded on many of the traits that belonged to the factors that made up the second-order Pro-sociality and Boldness domains, respectively. The second factor was an Impulsiveness factor. The correlation between the first and third factors was low, but the correlation between Impulsiveness and Pro-sociality was between medium and large, and therefore was consistent with the definitions of these factors.

We then extracted five factors and subjected them to a promax-rotation (see Table S13). The five factors resembled those from our initial factor analyses as did the interfactor correlations. The scree plot indicated that there were two factors (see Figure S13) as did both the parallel analysis and the BIC. The first higher-order factor was, when reflected, a Pro- sociality factor. The second higher-order factor was a Boldness factor (see Table S14).

#### Subjective Well-Being

Inspection of the scree plot (see Figure S14), parallel analysis, and the BIC all indicated that there was a single factor.^3^ This factor explained 67% of the variance and had salient loadings on all four items (see Table 5).

**Table 5.** Results from Factor Analysis of the Subjective Well-Being Scale

| Item | Loading | $h^2$ |
| --- | --- | --- |
| Be marmoset | 0.98 | 0.97 |
| Balance of moods | 0.86 | 0.73 |
| Ability to achieve goals | 0.82 | 0.67 |
| Pleasure from social interactions | 0.55 | 0.30 |

*Note.* $N = 128$ . $h^2$ = communalities.

### Reliabilities of Factors

#### Personality

The interrater reliabilities of the individual ratings for Sociability, Dominance, Impulsiveness, Openness, and Negative Affect were 0.52, 0.39, 0.28, 0.26, and 0.25, respectively. The interrater reliabilities of mean ratings for these factors were 0.74, 0.63, 0.50, 0.48, and 0.46, respectively. For Pro-sociality and Boldness, respectively, the interrater reliabilities of individual ratings were 0.45 and 0.30, and the interrater reliabilities of mean ratings for these second-order factors were 0.69 and 0.53.

The internal consistency reliability (Cronbach’s α) for Sociability, Dominance, Impulsiveness, Openness, and Negative Affect were 0.95, 0.95, 0.88, 0.85, and 0.81, respectively. The degree to which a general factor saturated these factors (McDonald’s ω*_h_*) was 0.81, 0.85, 0.75, 0.80, and 0.68, respectively.

#### Subjective Well-Being

For the total subjective well-being score, the interrater reliability of individual ratings was 0.21 and the interrater reliability of the mean of ratings 0.41. Cronbach’s α for this scale was 0.87 and McDonald’s ω*_h_* was 0.13.

### Personality Factor Comparisons

Iwanicki and Lehmann [17] found four factors. Compared to our first-order factors, their Extraversion factor overlapped with Dominance and Negative Affect, their Agreeableness factor overlapped with Sociability and (low) Dominance, their Conscientiousness factor overlapped with Sociability, and their Openness factor overlapped with the same-named factor that we found (see Table 6). Compared to our second-order factors, Iwanicki and Lehman’s Extraversion overlapped with (low) Pro-sociality and high Boldness; their Agreeableness and Conscientiousness factors overlapped with Pro-sociality; and their Openness factor overlapped with Boldness (see Table 7).

**Table 6.**
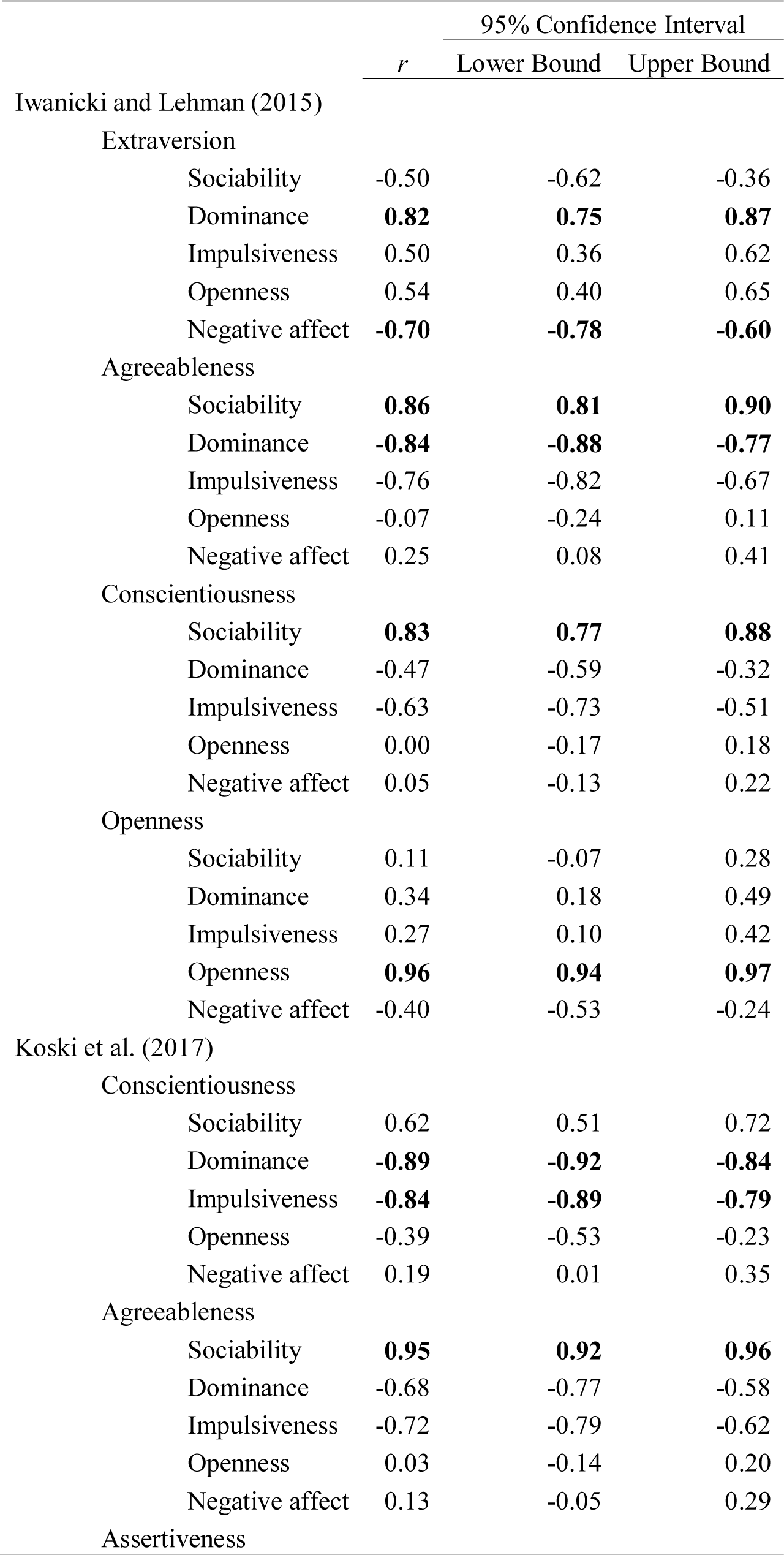
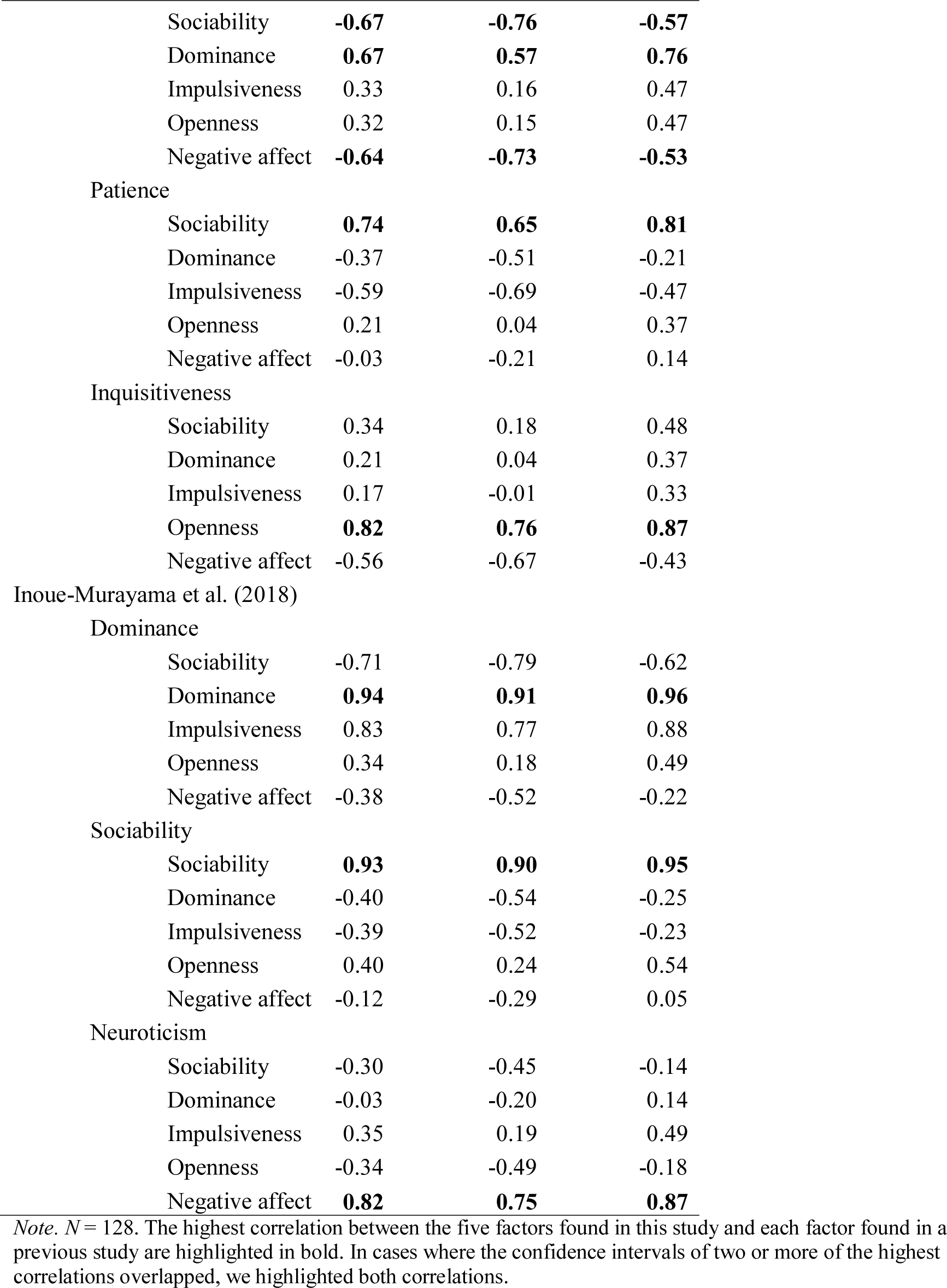
Correlations Between Unit-Weighted Factor Scores Based on Factor Loadings in the Present Study and Factor Loadings from Three Previous Studies of Common Marmoset Personality

|  |  | 95% Confidence Interval |  |  |
| --- | --- | --- | --- | --- |
|  |  | *r* | Lower Bound | Upper Bound |
| Iwanicki and Lehman (2015) |  |  |  |  |
| Extraversion |  |  |  |  |
|  | Sociability | -0.50 | -0.62 | -0.36 |
|  | Dominance | **0.82** | **0.75** | **0.87** |
|  | Impulsiveness | 0.50 | 0.36 | 0.62 |
|  | Openness | 0.54 | 0.40 | 0.65 |
|  | Negative affect | **-0.70** | **-0.78** | **-0.60** |
| Agreeableness |  |  |  |  |
|  | Sociability | **0.86** | **0.81** | **0.90** |
|  | Dominance | **-0.84** | **-0.88** | **-0.77** |
|  | Impulsiveness | -0.76 | -0.82 | -0.67 |
|  | Openness | -0.07 | -0.24 | 0.11 |
|  | Negative affect | 0.25 | 0.08 | 0.41 |
| Conscientiousness |  |  |  |  |
|  | Sociability | **0.83** | **0.77** | **0.88** |
|  | Dominance | -0.47 | -0.59 | -0.32 |
|  | Impulsiveness | -0.63 | -0.73 | -0.51 |
|  | Openness | 0.00 | -0.17 | 0.18 |
|  | Negative affect | 0.05 | -0.13 | 0.22 |
| Openness |  |  |  |  |
|  | Sociability | 0.11 | -0.07 | 0.28 |
|  | Dominance | 0.34 | 0.18 | 0.49 |
|  | Impulsiveness | 0.27 | 0.10 | 0.42 |
|  | Openness | **0.96** | **0.94** | **0.97** |
|  | Negative affect | -0.40 | -0.53 | -0.24 |
| Koski et al. (2017) |  |  |  |  |
| Conscientiousness |  |  |  |  |
|  | Sociability | 0.62 | 0.51 | 0.72 |
|  | Dominance | **-0.89** | **-0.92** | **-0.84** |
|  | Impulsiveness | **-0.84** | **-0.89** | **-0.79** |
|  | Openness | -0.39 | -0.53 | -0.23 |
|  | Negative affect | 0.19 | 0.01 | 0.35 |
| Agreeableness |  |  |  |  |
|  | Sociability | **0.95** | **0.92** | **0.96** |
|  | Dominance | -0.68 | -0.77 | -0.58 |
|  | Impulsiveness | -0.72 | -0.79 | -0.62 |
|  | Openness | 0.03 | -0.14 | 0.20 |
|  | Negative affect | 0.13 | -0.05 | 0.29 |
| Assertiveness |  |  |  |  |
|  | Sociability | **-0.67** | **-0.76** | **-0.57** |
|  | Dominance | **0.67** | **0.57** | **0.76** |
|  | Impulsiveness | 0.33 | 0.16 | 0.47 |
|  | Openness | 0.32 | 0.15 | 0.47 |
|  | Negative affect | **-0.64** | **-0.73** | **-0.53** |
| Patience |  |  |  |  |
|  | Sociability | **0.74** | **0.65** | **0.81** |
|  | Dominance | -0.37 | -0.51 | -0.21 |
|  | Impulsiveness | -0.59 | -0.69 | -0.47 |
|  | Openness | 0.21 | 0.04 | 0.37 |
|  | Negative affect | -0.03 | -0.21 | 0.14 |
| Inquisitiveness |  |  |  |  |
|  | Sociability | 0.34 | 0.18 | 0.48 |
|  | Dominance | 0.21 | 0.04 | 0.37 |
|  | Impulsiveness | 0.17 | -0.01 | 0.33 |
|  | Openness | **0.82** | **0.76** | **0.87** |
|  | Negative affect | -0.56 | -0.67 | -0.43 |
| Inoue-Murayama et al. (2018) |  |  |  |  |
| Dominance |  |  |  |  |
|  | Sociability | -0.71 | -0.79 | -0.62 |
|  | Dominance | **0.94** | **0.91** | **0.96** |
|  | Impulsiveness | 0.83 | 0.77 | 0.88 |
|  | Openness | 0.34 | 0.18 | 0.49 |
|  | Negative affect | -0.38 | -0.52 | -0.22 |
| Sociability |  |  |  |  |
|  | Sociability | **0.93** | **0.90** | **0.95** |
|  | Dominance | -0.40 | -0.54 | -0.25 |
|  | Impulsiveness | -0.39 | -0.52 | -0.23 |
|  | Openness | 0.40 | 0.24 | 0.54 |
|  | Negative affect | -0.12 | -0.29 | 0.05 |
| Neuroticism |  |  |  |  |
|  | Sociability | -0.30 | -0.45 | -0.14 |
|  | Dominance | -0.03 | -0.20 | 0.14 |
|  | Impulsiveness | 0.35 | 0.19 | 0.49 |
|  | Openness | -0.34 | -0.49 | -0.18 |
|  | Negative affect | **0.82** | **0.75** | **0.87** |

*Note.* $N = 128$ . The highest correlation between the five factors found in this study and each factor found in a previous study are highlighted in bold. In cases where the confidence intervals of two or more of the highest correlations overlapped, we highlighted both correlations.

**Table 7.**
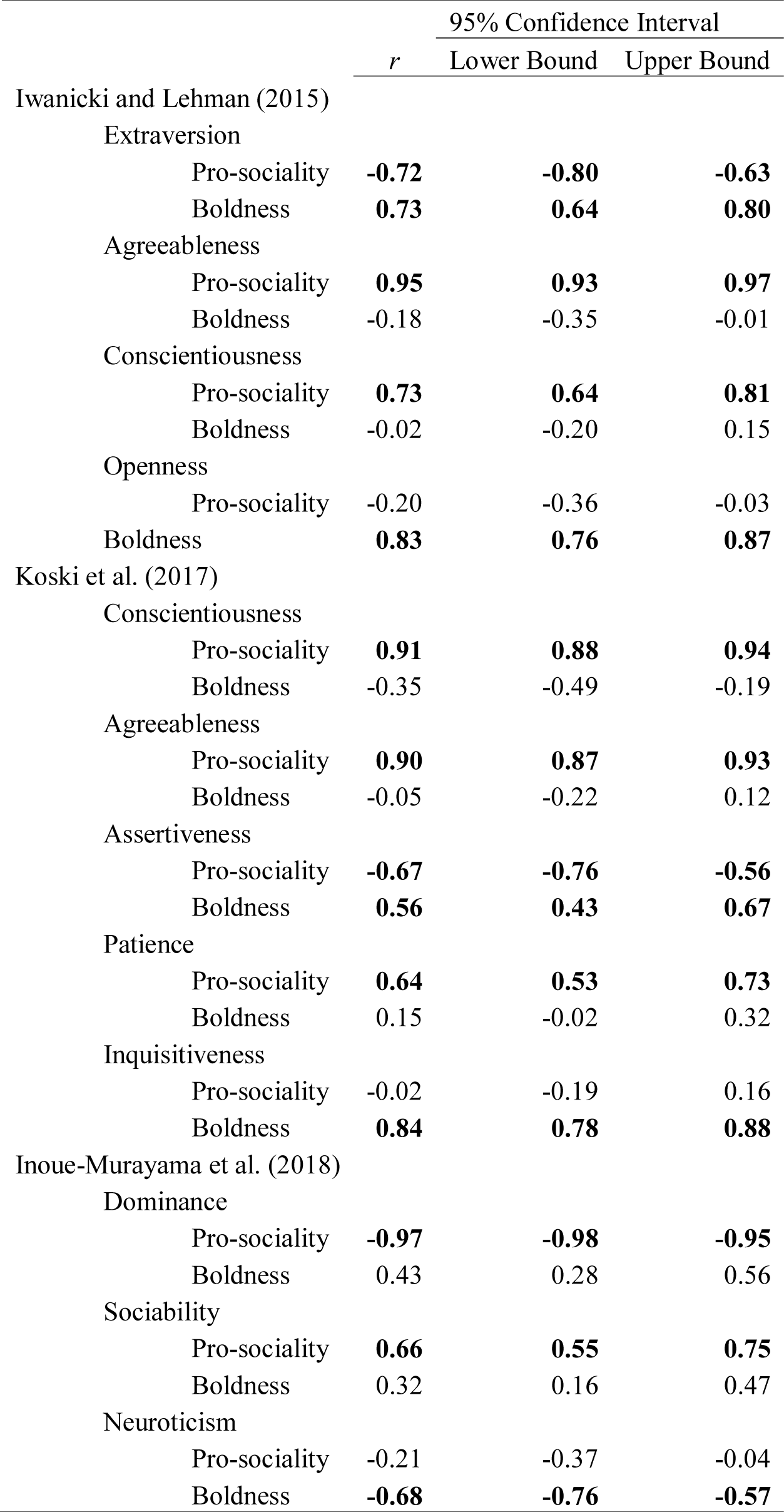
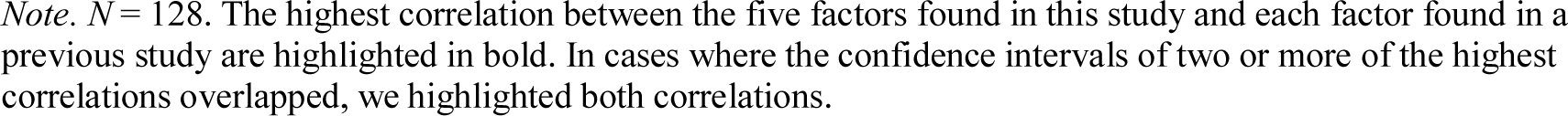
Correlations Between Unit-Weighted Factor Scores Based on Loadings from the Second-Order Factor Analysis in the Present Study and Factor Loadings from Three Previous Studies of Common Marmoset Personality

Koski et al. [31] found five factors. Compared to our first-order factors, their Conscientiousness factor overlapped with (low) Dominance and (low) Impulsiveness; their Agreeableness factor overlapped with Sociability; their Assertiveness factor overlapped with (low) Sociability, Dominance, and (low) Negative Affect; their Patience factor overlapped with Sociability; and their Inquisitiveness factor overlapped with Openness (see Table 6). Compared to our second-order factors, their Conscientiousness, Agreeableness, and Patience factors all overlapped with Pro-sociality; their Assertiveness factor overlapped with Boldness and (low) Pro-sociality; and their Inquisitiveness factor overlapped with Boldness (see Table 7).

Inoue-Murayama et al. [13] found three factors. Compared to our first-order factors, their Dominance factor overlapped with Dominance; their Sociability factor overlapped with Sociability; and their Neuroticism factor overlapped with Negative Affect (see Table 6). Compared to our second-order factors, their Dominance and Sociability factors overlapped with low and high Pro-sociality, respectively; their Neuroticism factor overlapped with (low) Boldness (see Table 7).

### Personality and Subjective Well-Being Associations

The correlations between the subjective well-being items and the personality factors are presented in Table 8. Sociability was significantly associated with higher scores on all four items and the total subjective well-being score. Dominance was not significantly related to any of the scale’s items or the factor. Impulsiveness was significantly related to lower, and Openness was significantly related to higher, balance of positive versus negative moods, how happy raters thought they would be if they were the marmoset, and the total subjective well-being score. Negative Affect was negatively related to how happy raters would be how happy raters thought they would be if they were the marmoset. Of the second-order factors, Pro- sociality was significantly associated with being rated as higher on all items save for the ability to achieve goals, and the total subjective well-being score. Boldness was not significantly associated with the pleasure subjects derived from social interactions, but it was significantly related to being higher in the other three items and in the total subjective well-being score.

**Table 8.** Correlations Between Subjective Well-Being Scale Items and Factor and Personality Factors

|  |  | 95% Confidence Interval |  |  |
| --- | --- | --- | --- | --- |
|  |  | *r* | Lower Bound | Upper Bound |
| Moods |  |  |  |  |
|  | Sociability | **0.42** | **0.26** | **0.55** |
|  | Dominance | -0.18 | -0.34 | < -0.01 |
|  | Impulsiveness | **-0.27** | **-0.43** | **-0.11** |
|  | Openness | **0.38** | **0.22** | **0.52** |
|  | Negative affect | -0.14 | -0.30 | 0.04 |
|  | Pro-sociality | **0.33** | **0.16** | **0.48** |
|  | Boldness | **0.31** | **0.15** | **0.46** |
| Social |  |  |  |  |
|  | Sociability | **0.53** | **0.40** | **0.65** |
|  | Dominance | -0.22 | -0.38 | -0.05 |
|  | Impulsiveness | -0.24 | -0.40 | -0.07 |
|  | Openness | 0.14 | -0.03 | 0.31 |
|  | Negative affect | -0.11 | -0.27 | 0.07 |
|  | Pro-sociality | **0.38** | **0.22** | **0.52** |
|  | Boldness | 0.15 | -0.03 | 0.31 |
| Goals |  |  |  |  |
|  | Sociability | **0.39** | **0.24** | **0.53** |
|  | Dominance | -0.08 | -0.25 | 0.10 |
|  | Impulsiveness | -0.22 | -0.38 | -0.05 |
|  | Openness | 0.25 | 0.08 | 0.40 |
|  | Negative affect | -0.22 | -0.38 | -0.05 |
|  | Pro-sociality | 0.26 | 0.09 | 0.41 |
|  | Boldness | **0.28** | **0.11** | **0.43** |
| Be marmoset |  |  |  |  |
|  | Sociability | **0.45** | **0.30** | **0.58** |
|  | Dominance | -0.18 | -0.34 | < -0.01 |
|  | Impulsiveness | **-0.28** | **-0.43** | **-0.11** |
|  | Openness | **0.33** | **0.16** | **0.48** |
|  | Negative affect | **-0.27** | **-0.42** | **-0.10** |
|  | Pro-sociality | **0.34** | **0.18** | **0.49** |
|  | Boldness | **0.36** | **0.20** | **0.50** |
| Subjective well-being |  |  |  |  |
|  | Sociability | **0.53** | **0.39** | **0.65** |
|  | Dominance | -0.19 | -0.36 | -0.02 |
|  | Impulsiveness | **-0.30** | **-0.45** | **-0.13** |
|  | Openness | **0.32** | **0.15** | **0.47** |
|  | Negative affect | -0.21 | -0.37 | -0.04 |
|  | Pro-sociality | **0.39** | **0.23** | **0.53** |
|  | Boldness | **0.32** | **0.16** | **0.47** |

*Note.* $N = 128$ . Estimates in bold are significant at $P < 0.05$ after adjustment using Holm's method. 95% confidence intervals are based on the nominal (unadjusted) significance level.

### Personality-Genotype Associations

#### G840C Genotypes

Twenty-seven subjects had the GG genotype, 23 had the CC genotype, and 72 were heterozygous. In the first set of analyses, we compared subjects with the GC genotype and subjects with the GG genotype to those with the CC genotype. Compared to subjects with the CC genotype, subjects with the GC or GG genotypes were significantly higher in Dominance; these associations, however, did not prevail correction for multiple tests (see Table S15). In a second set of analyses, we compared the 95 subjects who were carriers of the C allele (CC or GC genotype) to the 27 subjects with the GG genotype. None of the comparisons were statistically significant (see Table S16). In a third set of analyses, we compared the 99 subjects who were carriers of the G allele (GG or GC genotype) to the 23 subjects with the CC genotype. Carriers were significantly higher in Dominance, but this effect did not prevail correction for multiple tests (see Table S16).

#### T901A genotypes

Twelve subjects had the TT genotype, 35 had the AA genotype, and 76 were heterozygous. Because of this imbalance in the number of subjects, we only compared the 88 subjects who carried the T allele to the 35 subjects with the AA genotype. None of the comparisons were statistically significant (see Table S17).

## Discussion

We found five personality domains—Sociability, Dominance, Impulsiveness, Openness, and Negative Affect—in common marmosets. We also found two higher-order domains. One higher-order factor, Pro-sociality, had a positive loading on Sociability and negative loadings on Dominance and Impulsiveness, and a second, Boldness, which had a positive loading on Openness and a negative loading on Negative Affect. The interrater reliabilities of both sets of domains were comparable to what has been found in studies of other primate species [80], including humans [e.g., 81] and they were related to subjective well-being in ways consistent with their meaning. We found no strong evidence that either personality or subjective well-being was associated with the serotonin 1a receptor gene.

The personality domains that we found overlapped, although not completely, with those found in prior studies of common marmosets. Openness resembled eponymous domains, or a domain labeled Inquisitiveness, identified in previous studies [13, 17, 31, 32]. Moreover, although we did not find a clear Conscientiousness factor, as did two previous studies [17, 31], Impulsiveness and Pro-sociality overlapped with Conscientiousness in that all three were related to behavioral consistency and reliability, prosociality, tolerance, and low levels of aggression [17, 31]. Impulsiveness, however, was also related to emotionality and reactivity whereas Conscientiousness was not. Finally, Dominance, Sociability, and Negative Affect resembled domains found in earlier studies [13, 17, 31, 32]. On the other hand, although they may have been represented by Pro-sociality, we did not find strong evidence for a Patience [31] or a Perceptual Sensitivity [17] domain, although these may be the same construct [31] or related to Conscientiousness.

Unlike past rating-based studies that did not find higher-order factors in common marmosets [13, 17, 31], we found two such factors. Follow-on analyses indicated that these higher-order domains partly reflect a tendency for raters to see some traits as more correlated than others. However, these analyses could not exclude the possibility that these factors represented a higher-level of personality organization, perhaps reflecting group personalities [cf. 12]. Although there have been reports of higher-order factors of human personality [e.g., 82], including a so-called “general factor of personality” [e.g., 83], these reports have been criticized [e.g., 84, 85, 86]. The problems that affect human studies that purportedly find higher-order personality factors were absent in the present study: each animal was rated by two or three keepers, the correlations among latent variables were considerable, and adjusting for rater effects increased rather than decreased some interfactor correlations. Nevertheless, because second-order factors were not identified in other studies of common marmosets [13, 17, 31, 44], including one that included 77 of the same animals, prior to interpreting the meaning of this phenomenon, we urge an attempt to replicate these findings and an analysis of similar data using more flexible modeling techniques [e.g., 11].

The findings from the present study are consistent with the possibility that common marmosets evolved a personality structure that includes a domain or domains associated with self-control, such as the sort that are found in larger-brained primates, including brown capuchin monkeys [28], chimpanzees [39], and humans [59]. As these species share only a very distant common ancestors with Callitrichids and have very different socioecologies, these traits are not likely to be homologous. Instead, the presence of a domain related to self-control in common marmosets likely reflects convergent evolution that was driven by the need for individuals to meet the demands associated with cooperative breeding. Although, it is worth noting that there is still variability between studies in how these traits group, studies that examine the role that factors such as Conscientiousness, Patience, or Impulsivity play in infant rearing, especially by helpers, among common marmosets are needed to test this hypothesis.

As in our study of a subsample of these subjects [13], we found personality-subjective well-being correlations that were consistent with those found in studies of humans [45, 46] and nonhuman primates [40, 47-51]. These findings, and those in humans and great apes that indicate that a common genetic background underlies these traits [87-92], are consistent with the possibility that these relationships are ancestral.

Our failure to find association between SNPs related to the serotonin 1a receptor gene and either personality domains is not consistent with previous findings of an association between this genotype and personality in chimpanzees [54]. It is possible that our failure to find associations resulted from the personality measure that we used. However, as the associations between personality and serotonin-related genes in humans are likely false positives [93], we suspect that we did not find any associations because there were none.

This study had shortcomings. First, nearly 40% of the subjects were single housed. Behaviors related to some traits might therefore have been rare or absent, and so we still may not have been capturing enough between-subjects variation. Second, the factor structure was compared to studies that used different, although partly overlapping, instruments. It is unclear to what degree the use of different measures may have obscured similarities or blurred differences between the structures in these studies. This limitation also prevented us from using other statistical methods to directly compare these structures. Third, we judged that it was worth reporting the genetic associations so that they may contribute to future meta-analyses, as we alluded to previously, to identify genetic effects considerably larger sample sizes are needed. Fourth, the interrater reliabilities of the subjective well-being variables were lower than those reported in other nonhuman primate species, for example, chimpanzees [40, 47].

The cliché that a study’s findings can yield more questions than answers is well-suited for the present case. Nevertheless, these findings highlight the need for (and promise of) large collaborative studies if we are to understand the proximate and ultimate origins of personality structure in common marmosets, and other species, including ours.

## Data Availability

Data needed to reproduce the analyses are available via the Open Science Foundation website: https://osf.io/ysrja/.

## Footnotes

1 The HPQ can be obtained at https://extras.springer.com/2011/978-1-4614-0176-6.zip

2 We did not conduct a Hull test because the hullEFA function cannot be used to examine correlation matrices.

3 We conducted a Hull test for this analysis, too, but doing so produced a warning, which we suspect was attributable to there only being four items. The Hull test nevertheless indicated that there was one factor.

## Notes

This work was supported by Brain/MINDS Beyond from the Japan Agency of Medical Research and Development (AMED) (JP21dm0307006h0002) to TH, KAKENHI Grant Numbers 19H04904 and 20H00420 to MI-M, 18H05090 and 18K06372 to CY. We wish to thank the Leading graduate program in Primatology and Wildlife Science.

### Competing Interest Statement

The authors have declared no competing interest.

### Summary of Updates

In response to reviewer comments, we changed the figure into a table, which should be a clearer summary of the past literature. There were also other minor edits, such as a revised abstract and some edits for clarity. The author affiliations were changed a bit, namely the order of affiliations for AW were changed and one affiliation for MI-M was deleted.

