## Supplementary tables for "Personality, Subjective Well-Being, and the Serotonin 1a Receptor Gene in Common Marmosets (*Callithrix jacchus*)"

Table S1

*Definitions of Factors from Previous Studies*

| Study / Factor | Definition |
| --- | --- |
| Iwanicki and Lehman (2014) |  |
| Extraversion | dominant + stingy/greedy^a^ − submissive + independent − timid^b^ + defiant + reckless − depressed − cautious − solitary |
| Agreeableness | affectionate + helpful^c^ + cool^d^ + aggressive − irritable + gentle |
| Conscientiousness | predictable + protective + conventional + intelligent |
| Openness | active + curious + inventive |
| Koski et al. (2017) |  |
| Conscientiousness | − thoughtless − bullying − clumsy − reckless − disorganized − imitative − erratic − jealous − aggressive − irritable − impulsive − excitable − depressed − stingy/greedy^a^ − playful |
| Agreeableness | friendly + affectionate + gentle + sociable + helpful + predictable + unemotional + protective |
| Assertiveness | − cautious − dependent/follower^e^ + dominant + independent − timid − submissive − fearful − vulnerable − sympathetic |
| Patience | − distractible + intelligent + inventive + sensitive |
| Inquisitive | − lazy + inquisitive + active − solitary |
| Inoue-Murayama et al. (2018) |  |
| Dominance | defiant + stingy/greedy + jealous + aggressive + dominant + irritable + bullying + excitable + impulsive − submissive − friendly − gentle − cool + disorganized + erratic + active + manipulative − conventional − predictable + distractable + thoughtless |
| Sociability | helpful − solitary + imitative + dependent/follower + protective − individualistic − independent + sociable + sympathetic + affectionate + playful + sensitive + curious + inquisitive − lazy |
| Neuroticism | timid − stable + autistic + fearful + vulnerable − intelligent + clumsy + depressed |

*Note*. ^a^ Substituted for “+ stingy”. ^b^ Substituted for “+ bold”. ^c^ Substituted for “+ cooperative”. ^d^ Substituted for “+ cool”. ^e^ Substituted for “− dependent”.

Table S2

*Congruence Coefficients for Comparison of Varimax- and Promax-Rotated Structures*

|  | ML1 | ML4 | ML5 | ML2 | ML3 |
| --- | --- | --- | --- | --- | --- |
| ML1 | 0.95 | -0.37 | -0.32 | 0.12 | -0.08 |
| ML4 | -0.35 | 0.92 | 0.39 | 0.22 | -0.18 |
| ML5 | -0.32 | 0.40 | 0.94 | 0.24 | -0.06 |
| ML2 | 0.12 | 0.14 | 0.14 | 0.97 | -0.23 |
| ML3 | -0.05 | -0.11 | -0.03 | -0.19 | 0.97 |

Table S3

*Pattern Matrix from the First-Order Factor Analysis of the Hominoid Personality Questionnaire*

|  | Factor Loadings | | | | |  |
| --- | --- | --- | --- | --- | --- | --- |
| Item | Soc | Dom | Imp | Opn | Neg | *h*^2^ |
| Helpful | **0.85** | -0.10 | -0.16 | 0.12 | -0.13 | 0.79 |
| Sympathetic | **0.82** | -0.21 | -0.20 | 0.06 | 0.04 | 0.76 |
| Protective | **0.77** | -0.14 | -0.11 | 0.06 | -0.17 | 0.66 |
| Individualistic | **-0.77** | 0.22 | 0.11 | 0.07 | 0.23 | 0.71 |
| Sociable | **0.74** | **-0.40** | -0.31 | 0.15 | -0.17 | 0.85 |
| Dependent/follower | **0.73** | -0.15 | -0.14 | 0.16 | 0.31 | 0.69 |
| Solitary | **-0.71** | 0.14 | 0.08 | -0.24 | 0.38 | 0.74 |
| Independent | **-0.71** | 0.32 | -0.08 | 0.06 | 0.05 | 0.62 |
| Affectionate | **0.69** | -0.27 | -0.27 | 0.17 | 0.07 | 0.65 |
| Sensitive | **0.67** | -0.26 | -0.29 | -0.05 | 0.03 | 0.61 |
| Imitative | **0.66** | -0.10 | -0.03 | 0.18 | 0.11 | 0.49 |
| Friendly | **0.66** | **-0.56** | -0.32 | 0.08 | 0.03 | 0.86 |
| Gentle | **0.65** | **-0.50** | -0.38 | 0.08 | 0.08 | 0.83 |
| Conventional | **0.61** | -0.15 | -0.39 | -0.18 | 0.19 | 0.62 |
| Intelligent | **0.55** | 0.07 | -0.20 | 0.04 | -0.22 | 0.39 |
| Reckless | **-0.50** | 0.10 | **0.43** | **0.40** | -0.08 | 0.62 |
| Jealous | -0.24 | **0.82** | 0.20 | 0.25 | 0.01 | 0.83 |
| Stingy/greedy | -0.31 | **0.79** | 0.20 | 0.30 | 0.01 | 0.86 |
| Bullying | -0.29 | **0.78** | 0.23 | 0.16 | -0.03 | 0.77 |
| Dominant | -0.34 | **0.76** | 0.27 | 0.09 | -0.15 | 0.80 |
| Aggressive | -0.36 | **0.71** | 0.33 | 0.05 | -0.22 | 0.80 |
| Defiant | -0.32 | **0.69** | 0.36 | 0.10 | -0.25 | 0.78 |
| Manipulative | 0.03 | **0.62** | 0.06 | 0.14 | **-0.40** | 0.57 |
| Irritable | -0.27 | **0.58** | **0.55** | -0.07 | -0.12 | 0.74 |
| Excitable | -0.26 | 0.37 | **0.75** | 0.03 | -0.07 | 0.78 |
| Impulsive | -0.30 | 0.28 | **0.74** | 0.20 | 0.10 | 0.76 |
| Unemotional | 0.04 | -0.09 | **-0.66** | -0.13 | 0.25 | 0.52 |
| Cool | 0.35 | -0.28 | **-0.66** | -0.13 | 0.02 | 0.65 |
| Disorganized | -0.27 | 0.30 | **0.54** | 0.22 | 0.07 | 0.51 |
| Distractible | -0.21 | 0.13 | **0.51** | 0.26 | 0.06 | 0.40 |
| Stable | **0.40** | -0.30 | **-0.50** | 0.07 | -0.36 | 0.64 |
| Fearful | 0.05 | -0.02 | **0.47** | **-0.40** | 0.36 | 0.51 |
| Thoughtless | -0.28 | 0.15 | **0.43** | **0.40** | -0.05 | 0.45 |
| Erratic | -0.39 | **0.41** | **0.43** | -0.03 | 0.18 | 0.54 |
| Predictable | 0.24 | -0.21 | **-0.41** | -0.02 | 0.02 | 0.27 |
| Curious | 0.13 | 0.15 | 0.14 | **0.73** | -0.14 | 0.61 |
| Inquisitive | 0.16 | 0.12 | 0.11 | **0.70** | -0.07 | 0.55 |
| Playful | 0.22 | -0.02 | 0.28 | **0.67** | -0.16 | 0.60 |
| Inventive | 0.26 | 0.12 | 0.00 | **0.65** | -0.05 | 0.51 |
| Active | 0.16 | 0.27 | **0.42** | **0.60** | -0.25 | 0.69 |
| Cautious | 0.31 | -0.02 | 0.12 | **-0.57** | 0.30 | 0.53 |
| Timid | 0.00 | -0.01 | 0.28 | -0.25 | **0.66** | 0.58 |
| Autistic | 0.02 | -0.16 | 0.06 | 0.06 | **0.64** | 0.45 |
| Depressed | -0.16 | -0.04 | -0.21 | -0.19 | **0.64** | 0.51 |
| Vulnerable | 0.01 | -0.23 | -0.06 | -0.08 | **0.57** | 0.39 |
| Clumsy | -0.11 | 0.09 | 0.00 | -0.10 | **0.55** | 0.33 |
| Lazy | -0.14 | -0.18 | **-0.40** | -0.31 | **0.53** | 0.59 |
| Submissive | **0.40** | -0.35 | -0.28 | -0.12 | **0.49** | 0.61 |
| Proportion variance | 0.20 | 0.13 | 0.12 | 0.08 | 0.08 |  |

*Note*. *N* = 128. Soc = Sociability, Dom = Dominance, Imp = Impulsiveness, Opn = Openness, Neg = Negative Affect, *h*^2^ = communalities. Factors extracted using a maximum likelihood estimation and rotated using the varimax procedure. Factor loadings greater than or equal to |0.4| are in bold.

Table S4

*Pattern Matrix from the Factor Analysis of Residualized Ratings*

|  | Factor | |  |
| --- | --- | --- | --- |
| Item | I | II | *h*^2^ |
| Friendly | **-0.90** | 0.11 | 0.81 |
| Gentle | **-0.89** | 0.09 | 0.80 |
| Sociable | **-0.86** | 0.36 | 0.84 |
| Sympathetic | **-0.80** | 0.24 | 0.67 |
| Aggressive | **0.79** | 0.22 | 0.69 |
| Dominant | **0.77** | 0.22 | 0.67 |
| Stingy/greedy | **0.76** | 0.28 | 0.69 |
| Bullying | **0.75** | 0.20 | 0.63 |
| Affectionate | **-0.75** | 0.20 | 0.58 |
| Defiant | **0.74** | 0.30 | 0.67 |
| Helpful | **-0.74** | **0.42** | 0.69 |
| Sensitive | **-0.74** | 0.12 | 0.55 |
| Irritable | **0.73** | 0.16 | 0.57 |
| Jealous | **0.72** | 0.28 | 0.62 |
| Individualistic | **0.72** | -0.31 | 0.58 |
| Excitable | **0.70** | 0.19 | 0.54 |
| Protective | **-0.69** | 0.38 | 0.59 |
| Cool | **-0.68** | -0.16 | 0.50 |
| Conventional | **-0.68** | -0.12 | 0.48 |
| Erratic | **0.68** | -0.09 | 0.46 |
| Impulsive | **0.67** | 0.18 | 0.50 |
| Stable | **-0.67** | 0.24 | 0.48 |
| Independent | **0.66** | -0.19 | 0.46 |
| Disorganized | **0.61** | 0.19 | 0.43 |
| Dependent/follower | **-0.61** | 0.21 | 0.40 |
| Reckless | **0.58** | 0.25 | 0.42 |
| Submissive | **-0.57** | -0.35 | 0.48 |
| Intelligent | **-0.50** | 0.29 | 0.31 |
| Imitative | **-0.49** | 0.31 | 0.32 |
| Predictable | **-0.47** | -0.08 | 0.23 |
| Thoughtless | **0.45** | 0.29 | 0.30 |
| Distractible | **0.45** | 0.15 | 0.23 |
| Active | 0.23 | **0.74** | 0.63 |
| Lazy | -0.09 | **-0.70** | 0.51 |
| Curious | 0.07 | **0.64** | 0.42 |
| Playful | -0.05 | **0.64** | 0.40 |
| Solitary | **0.60** | **-0.62** | 0.69 |
| Inquisitive | 0.06 | **0.61** | 0.38 |
| Inventive | -0.05 | **0.61** | 0.37 |
| Depressed | 0.03 | **-0.59** | 0.35 |
| Timid | 0.15 | **-0.47** | 0.24 |
| Manipulative | 0.35 | **0.46** | 0.36 |
| Vulnerable | -0.11 | **-0.44** | 0.21 |
| Clumsy | 0.16 | -0.38 | 0.17 |
| Cautious | -0.15 | -0.37 | 0.17 |
| Unemotional | -0.29 | -0.37 | 0.23 |
| Autistic | 0.02 | -0.35 | 0.12 |
| Fearful | 0.17 | -0.32 | 0.12 |
| Proportion of variance | 0.34 | 0.13 |  |

*Note*. *N* = 128. Factors were not assigned labels. *h*^2^ = communalities. Factors extracted using a maximum likelihood estimation and rotated using the promax procedure. Factor loadings greater than or equal to |0.4| are in bold. Correlation between factors = 0.07.

Table S5

*Pattern Matrix from the Factor Analysis of Residualized Ratings*

|  | Factor | | | |  |
| --- | --- | --- | --- | --- | --- |
| Item | I | II | III | IV | *h*^2^ |
| Independent | **-0.83** | -0.10 | -0.06 | 0.07 | 0.63 |
| Helpful | **0.81** | -0.05 | -0.19 | 0.04 | 0.75 |
| Individualistic | **-0.80** | 0.02 | 0.22 | 0.12 | 0.69 |
| Dependent/follower | **0.78** | 0.09 | 0.23 | 0.14 | 0.61 |
| Sympathetic | **0.78** | -0.12 | 0.00 | 0.02 | 0.73 |
| Protective | **0.75** | -0.04 | -0.20 | 0.01 | 0.64 |
| Imitative | **0.71** | 0.15 | 0.07 | 0.13 | 0.45 |
| Solitary | **-0.70** | 0.04 | 0.34 | -0.19 | 0.71 |
| Sociable | **0.67** | -0.34 | -0.13 | 0.17 | 0.83 |
| Sensitive | **0.63** | -0.20 | -0.05 | -0.09 | 0.58 |
| Friendly | **0.59** | **-0.45** | 0.14 | 0.15 | 0.84 |
| Affectionate | **0.58** | -0.29 | 0.08 | 0.18 | 0.60 |
| Gentle | **0.55** | **-0.48** | 0.14 | 0.15 | 0.82 |
| Conventional | **0.49** | -0.24 | 0.02 | -0.26 | 0.52 |
| Intelligent | **0.42** | -0.17 | -0.34 | -0.06 | 0.37 |
| Excitable | 0.04 | **0.86** | 0.02 | 0.00 | 0.71 |
| Irritable | -0.07 | **0.80** | -0.19 | -0.18 | 0.71 |
| Impulsive | 0.00 | **0.79** | 0.22 | 0.20 | 0.72 |
| Cool | 0.09 | **-0.69** | -0.09 | -0.13 | 0.60 |
| Fearful | 0.32 | **0.63** | 0.39 | -0.38 | 0.51 |
| Stable | 0.20 | **-0.62** | -0.37 | 0.07 | 0.64 |
| Disorganized | -0.09 | **0.61** | 0.13 | 0.22 | 0.52 |
| Unemotional | -0.27 | **-0.59** | 0.02 | -0.16 | 0.34 |
| Erratic | -0.22 | **0.59** | 0.13 | -0.08 | 0.51 |
| Defiant | -0.25 | **0.58** | **-0.41** | -0.05 | 0.75 |
| Jealous | -0.24 | **0.56** | -0.27 | 0.04 | 0.64 |
| Aggressive | -0.33 | **0.56** | -0.38 | -0.09 | 0.76 |
| Dominant | -0.32 | **0.55** | -0.36 | -0.08 | 0.73 |
| Bullying | -0.31 | **0.54** | -0.27 | -0.02 | 0.65 |
| Stingy/greedy | -0.33 | **0.51** | -0.24 | 0.11 | 0.69 |
| Predictable | 0.06 | **-0.49** | -0.02 | 0.00 | 0.27 |
| Distractible | -0.08 | **0.42** | 0.23 | 0.30 | 0.35 |
| Autistic | -0.02 | 0.04 | **0.68** | 0.15 | 0.42 |
| Timid | 0.14 | **0.41** | **0.64** | -0.22 | 0.58 |
| Manipulative | -0.05 | 0.32 | **-0.63** | -0.05 | 0.53 |
| Vulnerable | -0.01 | -0.09 | **0.61** | 0.02 | 0.39 |
| Depressed | -0.25 | -0.16 | **0.51** | -0.14 | 0.40 |
| Submissive | 0.32 | -0.28 | **0.48** | -0.06 | 0.58 |
| Clumsy | -0.11 | 0.12 | **0.43** | -0.09 | 0.24 |
| Lazy | -0.33 | -0.37 | **0.42** | -0.26 | 0.52 |
| Playful | 0.26 | 0.11 | 0.04 | **0.72** | 0.60 |
| Curious | 0.09 | 0.06 | -0.06 | **0.71** | 0.57 |
| Inquisitive | 0.12 | 0.09 | -0.04 | **0.67** | 0.52 |
| Cautious | **0.45** | **0.40** | 0.15 | **-0.66** | 0.54 |
| Inventive | 0.19 | 0.05 | -0.09 | **0.60** | 0.45 |
| Reckless | -0.36 | 0.24 | 0.17 | **0.56** | 0.61 |
| Active | 0.27 | **0.45** | -0.17 | **0.54** | 0.68 |
| Thoughtless | -0.16 | 0.31 | 0.15 | **0.47** | 0.43 |
| Proportion of variance | 0.20 | 0.20 | 0.09 | 0.09 |  |
|  | Factor Correlations | | | |  |
|  | I | II | III | IV |  |
| I | 1.00 |  |  |  |  |
| II | -0.52 | 1.00 |  |  |  |
| III | 0.00 | -0.11 | 1.00 |  |  |
| IV | 0.02 | 0.25 | -0.33 | 1.00 |  |

*Note*. *N* = 128. Factors were not assigned labels. *h*^2^ = communalities. Factors extracted using a maximum likelihood estimation and rotated using the promax procedure. Factor loadings greater than or equal to |0.4| are in bold.

Table S6

*Pattern Matrix from the Factor Analysis of Residualized Ratings*

|  | Factor | | | | |  |
| --- | --- | --- | --- | --- | --- | --- |
| Item | Soc | Dom | Imp | Opn | Neg | *h*^2^ |
| Dependent/follower | **0.85** | 0.02 | 0.07 | 0.21 | 0.33 | 0.67 |
| Helpful | **0.85** | 0.07 | -0.09 | 0.10 | -0.08 | 0.77 |
| Sympathetic | **0.84** | 0.00 | -0.13 | 0.09 | 0.11 | 0.77 |
| Protective | **0.78** | 0.08 | -0.09 | 0.06 | -0.10 | 0.66 |
| Individualistic | **-0.70** | 0.13 | -0.08 | 0.15 | 0.29 | 0.70 |
| Imitative | **0.69** | -0.04 | 0.18 | 0.14 | 0.07 | 0.45 |
| Independent | **-0.68** | 0.31 | -0.31 | 0.14 | 0.12 | 0.66 |
| Sensitive | **0.64** | -0.08 | -0.16 | -0.05 | 0.00 | 0.59 |
| Affectionate | **0.63** | -0.10 | -0.24 | 0.25 | 0.18 | 0.65 |
| Conventional | **0.59** | 0.07 | -0.31 | -0.17 | 0.18 | 0.59 |
| Solitary | **-0.57** | 0.19 | -0.13 | -0.13 | **0.46** | 0.75 |
| Sociable | **0.57** | -0.35 | -0.10 | 0.13 | -0.22 | 0.84 |
| Intelligent | **0.54** | 0.27 | -0.35 | 0.04 | -0.12 | 0.43 |
| Gentle | **0.53** | -0.34 | -0.26 | 0.18 | 0.13 | 0.83 |
| Jealous | 0.02 | **0.84** | 0.02 | 0.16 | 0.07 | 0.77 |
| Dominant | -0.11 | **0.79** | 0.04 | 0.01 | -0.08 | 0.79 |
| Stingy/greedy | -0.09 | **0.77** | 0.01 | 0.24 | 0.08 | 0.80 |
| Bullying | -0.09 | **0.75** | 0.05 | 0.08 | 0.01 | 0.72 |
| Aggressive | -0.16 | **0.73** | 0.09 | -0.02 | -0.14 | 0.79 |
| Defiant | -0.08 | **0.73** | 0.12 | 0.02 | -0.17 | 0.77 |
| Manipulative | 0.12 | **0.68** | -0.11 | 0.04 | -0.36 | 0.56 |
| Irritable | 0.04 | **0.64** | 0.38 | -0.14 | -0.05 | 0.71 |
| Friendly | **0.45** | **-0.56** | -0.07 | 0.09 | -0.04 | 0.85 |
| Erratic | -0.09 | **0.47** | 0.27 | -0.03 | 0.26 | 0.54 |
| Impulsive | -0.03 | 0.18 | **0.68** | 0.14 | 0.13 | 0.72 |
| Cool | 0.19 | -0.06 | **-0.68** | -0.02 | 0.08 | 0.66 |
| Fearful | 0.21 | -0.07 | **0.66** | **-0.48** | 0.19 | 0.55 |
| Unemotional | -0.13 | 0.04 | **-0.66** | -0.05 | 0.23 | 0.43 |
| Excitable | 0.03 | 0.34 | **0.65** | -0.05 | -0.01 | 0.71 |
| Stable | 0.18 | -0.19 | **-0.48** | 0.09 | -0.34 | 0.64 |
| Distractible | -0.14 | -0.05 | **0.47** | 0.24 | 0.10 | 0.37 |
| Disorganized | -0.07 | 0.23 | **0.46** | 0.20 | 0.12 | 0.52 |
| Predictable | 0.12 | -0.09 | **-0.44** | 0.07 | 0.08 | 0.29 |
| Curious | 0.17 | 0.13 | -0.01 | **0.77** | 0.06 | 0.61 |
| Inquisitive | 0.17 | 0.08 | 0.07 | **0.70** | 0.03 | 0.53 |
| Playful | 0.23 | -0.12 | 0.22 | **0.69** | 0.00 | 0.59 |
| Inventive | 0.30 | 0.19 | -0.06 | **0.68** | 0.07 | 0.51 |
| Cautious | **0.44** | 0.14 | 0.27 | **-0.65** | 0.14 | 0.54 |
| Active | 0.25 | 0.16 | 0.39 | **0.51** | -0.18 | 0.68 |
| Reckless | **-0.41** | -0.09 | 0.32 | **0.50** | 0.06 | 0.62 |
| Thoughtless | -0.19 | -0.02 | 0.35 | **0.42** | 0.06 | 0.44 |
| Depressed | -0.10 | 0.06 | -0.25 | -0.03 | **0.68** | 0.52 |
| Autistic | 0.02 | -0.17 | 0.12 | 0.18 | **0.67** | 0.45 |
| Timid | 0.15 | -0.02 | 0.39 | -0.22 | **0.59** | 0.58 |
| Clumsy | 0.03 | 0.18 | -0.03 | 0.00 | **0.58** | 0.33 |
| Vulnerable | 0.01 | -0.23 | 0.02 | 0.04 | **0.58** | 0.40 |
| Submissive | 0.37 | -0.21 | -0.18 | 0.01 | **0.54** | 0.63 |
| Lazy | -0.24 | -0.09 | -0.36 | -0.18 | **0.51** | 0.57 |
| Proportion of variance | 0.18 | 0.15 | 0.12 | 0.09 | 0.08 |  |
|  | Factor Correlations | | | | |  |
|  | Soc | Dom | Imp | Opn | Neg |  |
| Soc | 1.00 |  |  |  |  |  |
| Dom | -0.56 | 1.00 |  |  |  |  |
| Imp | -0.40 | 0.50 | 1.00 |  |  |  |
| Opn | -0.02 | 0.21 | 0.23 | 1.00 |  |  |
| Neg | -0.17 | -0.12 | 0.07 | -0.38 | 1.00 |  |

*Note*. *N* = 128. Soc = Sociability, Dom = Dominance, Imp = Impulsiveness, Opn = Openness, Neg = Negative Affect, *h*^2^ = communalities. Factors extracted using a maximum likelihood estimation and rotated using the promax procedure. Factor loadings greater than or equal to |0.4| are in bold.

Table S7

*Congruence Coefficients for Comparison of Factors from Raw and Residualized Scores*

|  | MR5 | MR1 | MR4 | MR2 | MR3 |
| --- | --- | --- | --- | --- | --- |
| ML1 | 0.99 | -0.16 | -0.11 | 0.09 | -0.08 |
| ML4 | -0.12 | 0.98 | 0.12 | 0.10 | -0.07 |
| ML5 | -0.09 | 0.23 | 0.98 | 0.11 | -0.04 |
| ML2 | 0.15 | 0.07 | 0.06 | 0.99 | -0.02 |
| ML3 | 0.03 | -0.06 | 0.04 | -0.04 | 0.98 |

Table S8

*Pattern Matrix from the Factor Analysis of Rater 1’s Ratings*

|  | Factor | | | | |  |
| --- | --- | --- | --- | --- | --- | --- |
| Item | Imp | Soc | Dom | Opn | Neg | *h*^2^ |
| Impulsive | **0.94** | 0.06 | -0.07 | -0.01 | 0.05 | 0.77 |
| Excitable | **0.91** | 0.02 | -0.10 | -0.03 | 0.01 | 0.74 |
| Cool | **-0.74** | 0.01 | -0.08 | 0.04 | 0.03 | 0.63 |
| Erratic | **0.74** | 0.05 | 0.15 | -0.08 | -0.01 | 0.65 |
| Distractible | **0.69** | -0.21 | -0.07 | 0.20 | 0.05 | 0.62 |
| Stable | **-0.68** | -0.11 | -0.26 | 0.05 | -0.08 | 0.61 |
| Unemotional | **-0.67** | 0.10 | -0.02 | -0.06 | 0.27 | 0.67 |
| Disorganized | **0.65** | -0.19 | -0.01 | 0.09 | 0.07 | 0.58 |
| Predictable | **-0.64** | 0.21 | 0.15 | -0.28 | 0.03 | 0.53 |
| Reckless | **0.64** | -0.33 | -0.03 | 0.22 | 0.03 | 0.72 |
| Fearful | **0.60** | 0.26 | 0.02 | -0.30 | 0.25 | 0.47 |
| Irritable | **0.58** | 0.14 | 0.36 | -0.18 | -0.26 | 0.69 |
| Thoughtless | **0.57** | -0.19 | 0.02 | 0.28 | 0.06 | 0.50 |
| Timid | **0.54** | 0.08 | 0.16 | -0.06 | **0.42** | 0.51 |
| Clumsy | 0.32 | -0.02 | -0.11 | -0.09 | 0.30 | 0.20 |
| Protective | 0.20 | **0.87** | -0.13 | 0.07 | -0.35 | 0.77 |
| Helpful | 0.00 | **0.85** | -0.03 | 0.11 | -0.24 | 0.81 |
| Intelligent | -0.12 | **0.78** | 0.15 | -0.09 | -0.12 | 0.58 |
| Sympathetic | -0.03 | **0.76** | -0.15 | 0.03 | -0.17 | 0.75 |
| Affectionate | 0.00 | **0.63** | -0.19 | 0.18 | -0.12 | 0.65 |
| Sensitive | -0.24 | **0.61** | 0.04 | 0.13 | 0.24 | 0.72 |
| Dependent/follower | -0.02 | **0.57** | -0.07 | 0.39 | 0.20 | 0.65 |
| Independent | 0.13 | **-0.54** | 0.13 | 0.07 | -0.01 | 0.50 |
| Individualistic | 0.24 | **-0.48** | 0.23 | 0.15 | 0.18 | 0.61 |
| Imitative | -0.15 | **0.46** | 0.11 | 0.33 | 0.25 | 0.46 |
| Gentle | -0.28 | **0.45** | -0.29 | 0.13 | 0.08 | 0.83 |
| Cautious | 0.29 | **0.45** | -0.03 | -0.34 | 0.28 | 0.40 |
| Solitary | 0.21 | **-0.42** | 0.16 | -0.22 | 0.26 | 0.61 |
| Conventional | -0.38 | **0.41** | 0.03 | -0.10 | 0.36 | 0.67 |
| Sociable | -0.25 | **0.40** | -0.38 | 0.19 | -0.06 | 0.82 |
| Bullying | 0.02 | 0.01 | **0.91** | 0.05 | -0.02 | 0.84 |
| Aggressive | 0.03 | -0.05 | **0.86** | -0.01 | -0.11 | 0.88 |
| Jealous | -0.03 | -0.07 | **0.86** | 0.21 | 0.13 | 0.76 |
| Dominant | -0.01 | -0.05 | **0.85** | 0.03 | -0.09 | 0.80 |
| Stingy/greedy | 0.04 | -0.21 | **0.78** | 0.23 | 0.20 | 0.82 |
| Defiant | 0.07 | 0.02 | **0.78** | 0.02 | -0.13 | 0.70 |
| Manipulative | -0.17 | **0.47** | **0.51** | 0.23 | -0.21 | 0.42 |
| Friendly | -0.30 | 0.23 | **-0.42** | 0.25 | 0.22 | 0.83 |
| Curious | 0.09 | 0.10 | 0.07 | **0.80** | 0.07 | 0.61 |
| Inquisitive | -0.01 | 0.11 | 0.03 | **0.78** | 0.02 | 0.64 |
| Inventive | -0.04 | 0.14 | 0.12 | **0.76** | 0.15 | 0.57 |
| Playful | 0.16 | 0.12 | -0.05 | **0.76** | 0.01 | 0.58 |
| Active | 0.36 | 0.01 | 0.14 | **0.69** | -0.03 | 0.59 |
| Autistic | 0.06 | -0.09 | -0.07 | 0.39 | **0.66** | 0.41 |
| Vulnerable | 0.03 | -0.19 | -0.04 | 0.01 | **0.51** | 0.27 |
| Submissive | -0.14 | 0.31 | -0.14 | 0.06 | **0.47** | 0.54 |
| Lazy | -0.30 | -0.06 | -0.02 | -0.19 | **0.43** | 0.34 |
| Depressed | -0.18 | -0.21 | 0.27 | -0.12 | **0.43** | 0.29 |
| Proportion of variance | 0.19 | 0.16 | 0.13 | 0.09 | 0.05 |  |
|  | Factor Correlations | | | | |  |
|  | Imp | Soc | Dom | Opn | Neg |  |
| Imp | 1.00 |  |  |  |  |  |
| Soc | -0.62 | 1.00 |  |  |  |  |
| Dom | 0.51 | -0.55 | 1.00 |  |  |  |
| Opn | -0.15 | 0.18 | -0.08 | 1.00 |  |  |
| Neg | -0.04 | 0.15 | -0.17 | -0.34 | 1.00 |  |

*Note*. *N* = 128. Imp = Impulsiveness, Soc = Sociability, Dom = Dominance, Opn = Openness, Neg = Negative Affect, *h*^2^ = communalities. Factors extracted using a maximum likelihood estimation and rotated using the promax procedure. Factor loadings greater than or equal to |0.4| are in bold.

Table S9

*Pattern Matrix from Second-Order Factor Analysis from Rater 1’s Ratings*

|  | Second-order Factor | |  |
| --- | --- | --- | --- |
| First-order Factor | I | II | *h*^2^ |
| Sociability | **-0.83** | 0.03 | 0.68 |
| Impulsivity | **0.74** | 0.07 | 0.57 |
| Dominance | **0.68** | -0.07 | 0.45 |
| Negative affect | -0.29 | **< 1.00** | < 1.00 |
| Openness | -0.14 | -0.38 | 0.18 |
| Proportion of variance | 0.35 | 0.22 |  |

*Note*. *N* = 128. I = Pro-sociality (reversed), II = Boldness, *h*^2^ = communalities. ^1^ When rounded to three decimal places, this loading was equal to 0.995. Factors extracted using a maximum likelihood estimation and rotated using the promax procedure. Factor loadings greater than or equal to |0.4| are in bold. Factor correlation = -0.14.

Table S10

*Pattern Matrix from the Factor Analysis of Rater 2’s Ratings*

|  | Factor | | | |  |
| --- | --- | --- | --- | --- | --- |
| Item | I | II | III | IV | *h*^2^ |
| Aggressive | **0.81** | 0.30 | -0.19 | -0.16 | 0.64 |
| Irritable | **0.80** | -0.01 | 0.10 | -0.08 | 0.59 |
| Excitable | **0.79** | 0.17 | 0.08 | -0.09 | 0.56 |
| Dominant | **0.79** | 0.33 | -0.28 | -0.13 | 0.67 |
| Defiant | **0.75** | 0.24 | -0.09 | -0.16 | 0.51 |
| Gentle | **-0.73** | 0.31 | 0.07 | 0.12 | 0.71 |
| Manipulative | **0.72** | **0.40** | -0.13 | -0.10 | 0.54 |
| Submissive | **-0.70** | -0.13 | 0.03 | -0.02 | 0.49 |
| Affectionate | **-0.64** | 0.33 | 0.13 | 0.07 | 0.66 |
| Friendly | **-0.58** | 0.20 | 0.38 | 0.11 | 0.70 |
| Vulnerable | **-0.54** | -0.10 | -0.08 | -0.10 | 0.34 |
| Bullying | **0.53** | 0.01 | -0.17 | 0.03 | 0.36 |
| Sympathetic | **-0.52** | 0.18 | 0.35 | -0.04 | 0.59 |
| Stingy/greedy | **0.51** | -0.04 | -0.20 | 0.16 | 0.43 |
| Jealous | **0.48** | 0.01 | -0.16 | 0.08 | 0.31 |
| Lazy | **-0.48** | **-0.43** | -0.28 | -0.21 | 0.62 |
| Impulsive | **0.48** | -0.29 | 0.16 | 0.27 | 0.47 |
| Depressed | **-0.46** | -0.39 | -0.24 | -0.13 | 0.47 |
| Cool | **-0.45** | 0.39 | -0.21 | -0.30 | 0.52 |
| Helpful | **-0.40** | 0.30 | 0.38 | -0.03 | 0.59 |
| Predictable | -0.31 | 0.02 | -0.13 | 0.30 | 0.11 |
| Conventional | -0.31 | 0.25 | 0.13 | -0.25 | 0.33 |
| Intelligent | 0.20 | **0.73** | -0.07 | -0.01 | 0.49 |
| Timid | -0.38 | **-0.64** | 0.08 | -0.21 | 0.58 |
| Sensitive | -0.16 | **0.62** | 0.04 | -0.19 | 0.49 |
| Stable | -0.10 | **0.57** | 0.15 | -0.07 | 0.45 |
| Clumsy | -0.25 | **-0.54** | -0.08 | 0.07 | 0.32 |
| Sociable | -0.34 | **0.43** | 0.37 | 0.09 | 0.68 |
| Protective | -0.37 | **0.42** | 0.33 | 0.01 | 0.63 |
| Inventive | 0.05 | **0.41** | -0.01 | 0.16 | 0.21 |
| Erratic | 0.26 | -0.34 | 0.04 | -0.09 | 0.21 |
| Autistic | -0.15 | -0.30 | -0.06 | -0.02 | 0.12 |
| Disorganized | 0.20 | -0.27 | 0.18 | 0.19 | 0.16 |
| Independent | 0.07 | 0.10 | **-0.85** | 0.08 | 0.69 |
| Individualistic | 0.06 | -0.03 | **-0.84** | 0.08 | 0.75 |
| Imitative | -0.15 | -0.05 | **0.70** | -0.07 | 0.53 |
| Solitary | 0.05 | -0.28 | **-0.62** | -0.05 | 0.63 |
| Dependent/follower | **-0.42** | -0.12 | **0.49** | -0.12 | 0.48 |
| Unemotional | 0.02 | -0.06 | -0.31 | -0.11 | 0.14 |
| Reckless | -0.06 | -0.06 | -0.07 | **0.83** | 0.63 |
| Thoughtless | -0.01 | -0.01 | 0.00 | **0.74** | 0.54 |
| Curious | 0.10 | 0.29 | -0.06 | **0.72** | 0.70 |
| Playful | -0.02 | 0.20 | 0.17 | **0.66** | 0.60 |
| Cautious | 0.29 | 0.01 | 0.15 | **-0.65** | 0.35 |
| Inquisitive | 0.09 | 0.05 | 0.06 | **0.46** | 0.27 |
| Fearful | 0.13 | **-0.42** | 0.26 | **-0.44** | 0.35 |
| Active | 0.37 | 0.26 | 0.31 | **0.40** | 0.61 |
| Distractible | 0.04 | -0.07 | 0.04 | 0.38 | 0.15 |
| Proportion of variance | 0.19 | 0.10 | 0.10 | 0.09 |  |
|  | Factor Correlations | | | |  |
|  | I | II | III | IV |  |
| I | 1.00 |  |  |  |  |
| II | -0.17 | 1.00 |  |  |  |
| III | -0.19 | 0.41 | 1.00 |  |  |
| IV | 0.36 | 0.16 | 0.14 | 1.00 |  |

*Note*. *N* = 128. Factors were not assigned labels. *h*^2^ = communalities. Factors extracted using a maximum likelihood estimation and rotated using the promax procedure. Factor loadings greater than or equal to |0.4| are in bold.

Table S11

*Pattern Matrix from the Factor Analysis of Rater 3’s Ratings*

|  | Factor | | |  |
| --- | --- | --- | --- | --- |
| Item | I | II | III | *h*^2^ |
| Aggressive | **-0.89** | -0.08 | -0.05 | 0.761 |
| Gentle | **0.82** | 0.15 | -0.03 | 0.687 |
| Dominant | **-0.82** | 0.02 | 0.04 | 0.683 |
| Bullying | **-0.80** | 0.03 | 0.00 | 0.649 |
| Defiant | **-0.80** | 0.06 | -0.02 | 0.646 |
| Stingy/greedy | **-0.80** | 0.11 | -0.05 | 0.648 |
| Friendly | **0.78** | 0.19 | 0.08 | 0.596 |
| Affectionate | **0.72** | 0.19 | 0.07 | 0.509 |
| Irritable | **-0.71** | -0.05 | 0.20 | 0.605 |
| Submissive | **0.68** | -0.11 | 0.25 | 0.474 |
| Jealous | **-0.66** | 0.12 | 0.17 | 0.537 |
| Cautious | **0.58** | -0.19 | 0.01 | 0.388 |
| Sociable | **0.58** | 0.39 | -0.18 | 0.535 |
| Intelligent | **0.55** | 0.07 | 0.00 | 0.305 |
| Sympathetic | **0.55** | **0.49** | 0.14 | 0.475 |
| Reckless | **-0.52** | 0.02 | 0.17 | 0.341 |
| Excitable | **-0.52** | -0.01 | **0.48** | 0.613 |
| Manipulative | **-0.49** | 0.16 | -0.18 | 0.270 |
| Cool | **0.47** | -0.19 | **-0.46** | 0.579 |
| Protective | **0.47** | 0.36 | 0.00 | 0.319 |
| Autistic | **0.46** | -0.18 | 0.34 | 0.299 |
| Sensitive | **0.46** | 0.19 | 0.00 | 0.232 |
| Erratic | **-0.45** | -0.04 | 0.14 | 0.252 |
| Disorganized | **-0.43** | 0.09 | 0.08 | 0.223 |
| Conventional | **0.40** | 0.03 | -0.24 | 0.267 |
| Thoughtless | -0.39 | 0.25 | 0.20 | 0.309 |
| Solitary | -0.16 | **-0.70** | 0.14 | 0.525 |
| Active | -0.32 | **0.59** | 0.09 | 0.506 |
| Depressed | 0.24 | **-0.58** | 0.33 | 0.497 |
| Lazy | 0.21 | **-0.54** | -0.18 | 0.405 |
| Helpful | **0.46** | **0.52** | 0.04 | 0.433 |
| Imitative | 0.17 | **0.51** | 0.04 | 0.276 |
| Playful | 0.15 | **0.50** | 0.07 | 0.257 |
| Inquisitive | -0.15 | **0.45** | -0.15 | 0.251 |
| Independent | -0.33 | **-0.41** | -0.16 | 0.254 |
| Curious | -0.07 | 0.35 | 0.13 | 0.149 |
| Clumsy | 0.14 | -0.30 | 0.27 | 0.174 |
| Dependent/follower | 0.10 | 0.24 | 0.02 | 0.063 |
| Inventive | -0.10 | 0.18 | -0.01 | 0.044 |
| Timid | 0.20 | -0.12 | **0.72** | 0.506 |
| Stable | 0.06 | 0.04 | **-0.64** | 0.434 |
| Impulsive | -0.19 | 0.15 | **0.62** | 0.496 |
| Unemotional | 0.32 | -0.25 | **-0.61** | 0.642 |
| Vulnerable | 0.35 | -0.22 | **0.57** | 0.422 |
| Predictable | 0.30 | -0.08 | **-0.47** | 0.390 |
| Fearful | 0.05 | 0.00 | **0.42** | 0.169 |
| Distractible | -0.08 | 0.23 | 0.39 | 0.225 |
| Individualistic | -0.22 | -0.14 | 0.30 | 0.190 |
| Proportion of variance | 0.24 | 0.08 | 0.08 |  |
|  | Factor Correlations | | |  |
|  | I | II | III |  |
| I | 1.00 | -0.08 | -0.24 |  |
| II | -0.08 | 1.00 | -0.02 |  |
| III | -0.24 | -0.02 | 1.00 |  |

*Note*. *N* = 81. Factors were not assigned labels. *h*^2^ = communalities. Factors extracted using a maximum likelihood estimation and rotated using the promax procedure. Factor loadings greater than or equal to |0.4| are in bold.

Table S12

*Pattern Matrix from the Factor Analysis of Weighted Correlation Matrix (***Ρ**_w_*)*

|  | Factor | | |  |
| --- | --- | --- | --- | --- |
| Item | Pro-sociality | Impulsiveness | Boldness | *h*^2^ |
| Friendly | **0.82** | -0.06 | -0.01 | 0.71 |
| Aggressive | **-0.79** | 0.00 | **0.40** | 0.72 |
| Dominant | **-0.78** | -0.02 | **0.42** | 0.69 |
| Gentle | **0.75** | -0.19 | -0.01 | 0.71 |
| Sociable | **0.73** | -0.10 | 0.26 | 0.70 |
| Bullying | **-0.70** | 0.04 | 0.32 | 0.57 |
| Affectionate | **0.70** | -0.08 | 0.06 | 0.56 |
| Sympathetic | **0.70** | -0.05 | 0.11 | 0.55 |
| Stingy/greedy | **-0.66** | 0.13 | 0.29 | 0.58 |
| Defiant | **-0.66** | 0.06 | 0.39 | 0.57 |
| Helpful | **0.65** | -0.05 | 0.26 | 0.56 |
| Protective | **0.64** | -0.01 | 0.25 | 0.51 |
| Jealous | **-0.59** | 0.12 | 0.30 | 0.48 |
| Independent | **-0.58** | -0.06 | -0.10 | 0.33 |
| Dependent/follower | **0.56** | 0.03 | 0.05 | 0.31 |
| Solitary | **-0.55** | -0.03 | **-0.45** | 0.55 |
| Irritable | **-0.54** | 0.30 | 0.20 | 0.55 |
| Submissive | **0.54** | -0.09 | -0.35 | 0.42 |
| Imitative | **0.51** | 0.08 | 0.20 | 0.30 |
| Individualistic | **-0.50** | 0.14 | -0.20 | 0.39 |
| Sensitive | **0.45** | -0.25 | 0.20 | 0.41 |
| Impulsive | -0.12 | **0.68** | -0.09 | 0.54 |
| Reckless | -0.11 | **0.63** | 0.06 | 0.47 |
| Cool | 0.23 | **-0.61** | 0.01 | 0.55 |
| Thoughtless | -0.01 | **0.61** | 0.13 | 0.41 |
| Distractible | 0.02 | **0.60** | -0.03 | 0.34 |
| Unemotional | -0.04 | **-0.59** | -0.04 | 0.33 |
| Disorganized | -0.16 | **0.47** | -0.05 | 0.31 |
| Excitable | -0.33 | **0.47** | 0.08 | 0.48 |
| Stable | 0.23 | **-0.45** | 0.33 | 0.44 |
| Predictable | 0.10 | **-0.45** | 0.07 | 0.24 |
| Conventional | 0.35 | -0.37 | -0.02 | 0.38 |
| Erratic | -0.31 | 0.36 | -0.15 | 0.35 |
| Manipulative | -0.37 | -0.21 | **0.59** | 0.39 |
| Timid | 0.06 | 0.33 | **-0.58** | 0.38 |
| Active | 0.10 | **0.51** | **0.52** | 0.56 |
| Depressed | -0.12 | -0.12 | **-0.51** | 0.31 |
| Curious | 0.18 | 0.31 | **0.49** | 0.37 |
| Lazy | -0.04 | -0.36 | **-0.49** | 0.40 |
| Vulnerable | 0.22 | 0.10 | **-0.45** | 0.21 |
| Playful | 0.39 | **0.40** | **0.43** | 0.44 |
| Clumsy | -0.01 | 0.12 | **-0.42** | 0.18 |
| Inquisitive | 0.19 | 0.22 | **0.42** | 0.27 |
| Fearful | -0.02 | 0.23 | -0.38 | 0.19 |
| Inventive | 0.20 | 0.13 | 0.38 | 0.21 |
| Autistic | 0.17 | 0.09 | -0.33 | 0.12 |
| Intelligent | 0.27 | -0.23 | 0.32 | 0.29 |
| Cautious | 0.05 | -0.20 | -0.25 | 0.12 |
| Proportion of variance | 0.22 | 0.11 | 0.10 |  |
|  | Factor Correlations | | |  |
|  | Pro-sociality | Impulsiveness | Boldness |  |
| Pro-sociality | 1.00 |  |  |  |
| Impulsiveness | -0.44 | 1.00 |  |  |
| Boldness | 0.11 | 0.11 | 1.00 |  |

*Note*. *N* = 128. *h*^2^ = communalities. Factors extracted using a maximum likelihood estimation and rotated using the promax procedure. Factor loadings greater than or equal to |0.4| are in bold.

Table S13

*Pattern Matrix from the Factor Analysis of the Weighted Correlation Matrix (***R**_w_*)*

|  | Factor | | | | |  |
| --- | --- | --- | --- | --- | --- | --- |
| Item | Soc | Imp | Dom | Opn | Neg | *h*^2^ |
| Helpful | **0.85** | 0.02 | 0.08 | 0.01 | 0.00 | 0.66 |
| Sympathetic | **0.78** | 0.01 | -0.03 | 0.01 | 0.10 | 0.62 |
| Protective | **0.76** | 0.05 | 0.00 | 0.02 | -0.05 | 0.57 |
| Affectionate | **0.65** | -0.12 | -0.07 | 0.13 | 0.18 | 0.61 |
| Dependent/follower | **0.63** | 0.03 | 0.03 | 0.08 | 0.21 | 0.38 |
| Independent | **-0.63** | -0.21 | 0.19 | 0.15 | 0.16 | 0.42 |
| Solitary | **-0.62** | -0.08 | 0.07 | -0.10 | 0.33 | 0.59 |
| Sociable | **0.60** | -0.13 | -0.18 | 0.16 | -0.11 | 0.70 |
| Imitative | **0.59** | 0.09 | 0.01 | 0.10 | 0.02 | 0.32 |
| Sensitive | **0.57** | -0.24 | 0.10 | 0.03 | 0.05 | 0.44 |
| Gentle | **0.56** | -0.27 | -0.16 | 0.15 | 0.22 | 0.75 |
| Friendly | **0.56** | -0.09 | -0.32 | 0.11 | 0.08 | 0.70 |
| Individualistic | **-0.54** | 0.00 | 0.17 | 0.18 | 0.31 | 0.48 |
| Conventional | **0.50** | -0.32 | 0.13 | -0.16 | 0.17 | 0.46 |
| Intelligent | **0.43** | -0.20 | 0.13 | 0.04 | -0.12 | 0.29 |
| Impulsive | -0.02 | **0.75** | -0.03 | 0.00 | 0.03 | 0.56 |
| Cool | 0.09 | **-0.71** | -0.01 | 0.03 | 0.09 | 0.58 |
| Unemotional | -0.19 | **-0.67** | 0.00 | -0.02 | 0.01 | 0.37 |
| Excitable | 0.03 | **0.62** | 0.22 | -0.13 | -0.09 | 0.55 |
| Distractible | -0.03 | **0.54** | -0.05 | 0.22 | 0.11 | 0.34 |
| Predictable | 0.02 | **-0.51** | 0.03 | 0.04 | 0.01 | 0.26 |
| Fearful | 0.22 | **0.49** | 0.01 | **-0.47** | 0.18 | 0.40 |
| Stable | 0.11 | **-0.48** | -0.11 | 0.05 | -0.37 | 0.50 |
| Thoughtless | -0.13 | **0.47** | -0.05 | **0.43** | 0.05 | 0.47 |
| Reckless | -0.29 | **0.47** | -0.09 | **0.45** | 0.08 | 0.56 |
| Irritable | -0.04 | **0.47** | **0.40** | -0.18 | -0.17 | 0.63 |
| Disorganized | -0.12 | **0.46** | 0.01 | 0.09 | 0.05 | 0.31 |
| Active | 0.23 | **0.44** | 0.10 | 0.39 | -0.24 | 0.56 |
| Erratic | -0.16 | **0.44** | 0.09 | -0.13 | 0.05 | 0.36 |
| Jealous | 0.00 | 0.06 | **0.79** | 0.17 | 0.23 | 0.66 |
| Dominant | -0.16 | 0.01 | **0.74** | 0.02 | -0.08 | 0.73 |
| Bullying | -0.12 | 0.03 | **0.73** | 0.06 | 0.06 | 0.65 |
| Stingy/greedy | -0.17 | 0.04 | **0.71** | 0.20 | 0.16 | 0.70 |
| Aggressive | -0.15 | 0.07 | **0.70** | -0.04 | -0.13 | 0.74 |
| Manipulative | 0.18 | -0.18 | **0.61** | 0.05 | -0.22 | 0.40 |
| Defiant | -0.13 | 0.12 | **0.56** | -0.02 | -0.17 | 0.58 |
| Curious | 0.18 | 0.06 | 0.20 | **0.69** | 0.08 | 0.54 |
| Playful | 0.28 | 0.21 | -0.03 | **0.62** | -0.01 | 0.53 |
| Inquisitive | 0.12 | 0.04 | 0.05 | **0.53** | -0.06 | 0.35 |
| Cautious | 0.34 | 0.04 | 0.11 | **-0.52** | 0.10 | 0.32 |
| Inventive | 0.18 | -0.05 | 0.13 | **0.49** | 0.05 | 0.28 |
| Vulnerable | 0.14 | 0.05 | -0.01 | 0.01 | **0.60** | 0.35 |
| Depressed | -0.17 | -0.17 | 0.08 | -0.10 | **0.57** | 0.43 |
| Timid | 0.09 | **0.41** | -0.03 | -0.21 | **0.55** | 0.53 |
| Autistic | 0.04 | -0.01 | -0.02 | 0.12 | **0.51** | 0.23 |
| Lazy | -0.19 | **-0.43** | -0.01 | -0.13 | **0.47** | 0.48 |
| Submissive | 0.37 | -0.12 | -0.15 | -0.02 | **0.45** | 0.51 |
| Clumsy | -0.07 | 0.12 | -0.03 | -0.10 | 0.38 | 0.22 |
| Proportion of variance | 0.16 | 0.12 | 0.10 | 0.06 | 0.06 |  |
|  | Factor Correlations | | | | |  |
|  | Soc | Imp | Dom | Opn | Neg |  |
| Soc | 1.00 |  |  |  |  |  |
| Imp | -0.51 | 1.00 |  |  |  |  |
| Dom | -0.49 | 0.48 | 1.00 |  |  |  |
| Opn | 0.15 | 0.10 | 0.09 | 1.00 |  |  |
| Neg | -0.10 | -0.03 | -0.26 | -0.39 | 1.00 |  |

*Note*. *N* = 128. Soc = Sociability, Dom = Dominance, Imp = Impulsiveness, Opn = Openness, Neg = Negative Affect, *h*^2^ = communalities. Factors extracted using a maximum likelihood estimation and rotated using the promax procedure. Factor loadings greater than or equal to |0.4| are in bold.

Table S14

*Pattern Matrix from Second-Order Factor Analysis from the Weighted Correlation Matrix (***R**_w_*)*

|  | Second-order factor | |  |
| --- | --- | --- | --- |
| First-order factor | Pro-sociality | Boldness | *h*^2^ |
| Sociability | **0.79** | 0.25 | 0.66 |
| Dominance | **-0.70** | 0.24 | 0.57 |
| Impulsiveness | **-0.66** | 0.01 | 0.44 |
| Negative affect | 0.04 | **-0.75** | 0.56 |
| Openness | 0.03 | **0.53** | 0.28 |
| Proportion of variance | 0.31 | 0.19 |  |

*Note*. *N* = 128. *h*^2^ = communalities. Factors extracted using a maximum likelihood estimation and rotated using the promax procedure. Factor loadings greater than or equal to |0.4| are in bold. Factor correlation = -0.08.

Table S15

*Effects of G840C Genotype on Personality Domains*

|  | *b* | *SE* | *t* | *P* |
| --- | --- | --- | --- | --- |
| Sociability |  |  |  |  |
| Intercept | 0.85 | 0.30 | 2.81 | 0.006 |
| Male vs. Female | -0.75 | 0.22 | -3.49 | < 0.001 |
| Age | -0.01 | 0.03 | -0.34 | 0.74 |
| GC vs. CC | -0.22 | 0.23 | -0.95 | 0.35 |
| GG vs. CC | -0.33 | 0.27 | -1.21 | 0.23 |
| Dominance |  |  |  |  |
| Intercept | -0.94 | 0.30 | -3.08 | 0.003 |
| Male vs. Female | 0.70 | 0.22 | 3.23 | 0.002 |
| Age | -0.01 | 0.03 | -0.20 | 0.84 |
| GC vs. CC | 0.51 | 0.23 | 2.19 | 0.03 |
| GG vs. CC | 0.58 | 0.27 | 2.12 | 0.036 |
| Impulsiveness |  |  |  |  |
| Intercept | -0.59 | 0.30 | -1.96 | 0.052 |
| Male vs. Female | 0.65 | 0.22 | 3.01 | 0.003 |
| Age | -0.02 | 0.03 | -0.68 | 0.50 |
| GC vs. CC | 0.17 | 0.23 | 0.73 | 0.47 |
| GG vs. CC | 0.37 | 0.27 | 1.35 | 0.18 |
| Openness |  |  |  |  |
| Intercept | -0.08 | 0.30 | -0.26 | 0.80 |
| Male vs. Female | 0.45 | 0.22 | 2.08 | 0.04 |
| Age | -0.05 | 0.03 | -1.41 | 0.16 |
| GC vs. CC | -0.03 | 0.23 | -0.12 | 0.91 |
| GG vs. CC | -0.09 | 0.27 | -0.31 | 0.75 |
| Negative Affect |  |  |  |  |
| Intercept | 0.48 | 0.32 | 1.50 | 0.14 |
| Male vs. Female | -0.14 | 0.23 | -0.63 | 0.53 |
| Age | -0.06 | 0.03 | -1.60 | 0.11 |
| GC vs. CC | -0.10 | 0.24 | -0.40 | 0.69 |
| GG vs. CC | -0.16 | 0.29 | -0.55 | 0.58 |
| Pro-sociality |  |  |  |  |
| Intercept | 0.94 | 0.30 | 3.16 | 0.002 |
| Male vs. Female | -0.81 | 0.21 | -3.83 | < 0.001 |
| Age | 0.01 | 0.03 | 0.18 | 0.86 |
| GC vs. CC | -0.37 | 0.23 | -1.63 | 0.11 |
| GG vs. CC | -0.51 | 0.27 | -1.90 | 0.06 |
| Boldness |  |  |  |  |
| Intercept | -0.32 | 0.31 | -1.01 | 0.31 |
| Male vs. Female | 0.36 | 0.22 | 1.63 | 0.11 |
| Age | 0.00 | 0.03 | 0.05 | 0.96 |
| GC vs. CC | 0.04 | 0.24 | 0.16 | 0.88 |
| GG vs. CC | 0.03 | 0.28 | 0.12 | 0.90 |

*Note*. *N* = 122.

Table S16

*Effects of G840C Genotype on Personality Domains*

|  | Genotype | | | | | | | | |
| --- | --- | --- | --- | --- | --- | --- | --- | --- | --- |
|  | C Present vs. Absent | | | |  | G Present vs. Absent | | | |
|  | *b* | *SE* | *t* | *P* |  | *b* | *SE* | *t* | *P* |
| Sociability |  |  |  |  |  |  |  |  |  |
| Intercept | 0.49 | 0.28 | 1.76 | 0.082 |  | 0.86 | 0.30 | 2.87 | 0.005 |
| Male vs. Female | -0.74 | 0.21 | -3.43 | < 0.001 |  | -0.75 | 0.21 | -3.50 | < 0.001 |
| Age | -0.01 | 0.03 | -0.23 | 0.82 |  | -0.01 | 0.03 | -0.40 | 0.69 |
| Genotype | 0.17 | 0.21 | 0.79 | 0.43 |  | -0.25 | 0.22 | -1.13 | 0.26 |
| Impulsiveness |  |  |  |  |  |  |  |  |  |
| Intercept | -0.28 | 0.29 | -1.00 | 0.32 |  | -0.94 | 0.30 | -3.12 | 0.002 |
| Male vs. Female | 0.66 | 0.22 | 3.02 | 0.003 |  | 0.70 | 0.22 | 3.24 | 0.002 |
| Age | -0.01 | 0.03 | -0.44 | 0.66 |  | -0.01 | 0.03 | -0.16 | 0.87 |
| Genotype | -0.20 | 0.21 | -0.93 | 0.36 |  | 0.53 | 0.22 | 2.37 | 0.019 |
| Dominance |  |  |  |  |  |  |  |  |  |
| Intercept | -0.20 | 0.28 | -0.72 | 0.47 |  | -0.61 | 0.30 | -2.03 | 0.044 |
| Male vs. Female | 0.64 | 0.21 | 2.97 | 0.004 |  | 0.65 | 0.22 | 3.01 | 0.003 |
| Age | -0.03 | 0.03 | -0.77 | 0.44 |  | -0.02 | 0.03 | -0.57 | 0.57 |
| Genotype | -0.24 | 0.21 | -1.16 | 0.25 |  | 0.23 | 0.22 | 1.01 | 0.31 |
| Openness |  |  |  |  |  |  |  |  |  |
| Intercept | -0.17 | 0.28 | -0.60 | 0.55 |  | -0.07 | 0.30 | -0.24 | 0.81 |
| Male vs. Female | 0.45 | 0.21 | 2.10 | 0.038 |  | 0.45 | 0.22 | 2.09 | 0.039 |
| Age | -0.05 | 0.03 | -1.41 | 0.16 |  | -0.05 | 0.03 | -1.46 | 0.15 |
| Genotype | 0.07 | 0.21 | 0.31 | 0.76 |  | -0.04 | 0.22 | -0.20 | 0.84 |
| Negative Affect |  |  |  |  |  |  |  |  |  |
| Intercept | 0.31 | 0.29 | 1.04 | 0.30 |  | 0.48 | 0.32 | 1.53 | 0.13 |
| Male vs. Female | -0.14 | 0.23 | -0.60 | 0.55 |  | -0.14 | 0.23 | -0.63 | 0.53 |
| Age | -0.05 | 0.03 | -1.57 | 0.12 |  | -0.06 | 0.03 | -1.65 | 0.10 |
| Genotype | 0.09 | 0.22 | 0.39 | 0.70 |  | -0.12 | 0.23 | -0.49 | 0.62 |
| Pro-sociality |  |  |  |  |  |  |  |  |  |
| Intercept | 0.38 | 0.28 | 1.38 | 0.17 |  | 0.96 | 0.30 | 3.22 | 0.002 |
| Male vs. Female | -0.79 | 0.21 | -3.69 | < 0.001 |  | -0.81 | 0.21 | -3.84 | < 0.001 |
| Age | 0.01 | 0.03 | 0.37 | 0.72 |  | 0.00 | 0.03 | 0.10 | 0.92 |
| Genotype | 0.23 | 0.21 | 1.11 | 0.27 |  | -0.41 | 0.22 | -1.87 | 0.064 |
| Boldness |  |  |  |  |  |  |  |  |  |
| Intercept | -0.28 | 0.29 | -0.96 | 0.34 |  | -0.32 | 0.31 | -1.02 | 0.31 |
| Male vs. Female | 0.36 | 0.22 | 1.63 | 0.11 |  | 0.36 | 0.22 | 1.64 | 0.10 |
| Age | 0.00 | 0.03 | 0.03 | 0.97 |  | 0.00 | 0.03 | 0.05 | 0.96 |
| Genotype | -0.01 | 0.22 | -0.03 | 0.98 |  | 0.04 | 0.23 | 0.16 | 0.87 |

*Note*. *N* = 122.

Table S17

*Effects of T901A Genotype on Personality Domains*

|  | *b* | *SE* | *t* | *P* |
| --- | --- | --- | --- | --- |
| Sociability |  |  |  |  |
| Intercept | 0.73 | 0.24 | 3.12 | 0.002 |
| Male vs. Female | -0.72 | 0.21 | -3.43 | < 0.001 |
| Age | 0.01 | 0.03 | 0.17 | 0.87 |
| T allele present | -0.26 | 0.19 | -1.39 | 0.17 |
| Impulsiveness |  |  |  |  |
| Intercept | -0.59 | 0.23 | -2.50 | 0.014 |
| Male vs. Female | 0.64 | 0.21 | 3.07 | 0.003 |
| Age | -0.03 | 0.03 | -0.89 | 0.38 |
| T allele present | 0.29 | 0.19 | 1.54 | 0.13 |
| Dominance |  |  |  |  |
| Intercept | -0.37 | 0.23 | -1.58 | 0.12 |
| Male vs. Female | 0.64 | 0.21 | 3.07 | 0.003 |
| Age | -0.04 | 0.03 | -1.19 | 0.24 |
| T allele present | 0.05 | 0.19 | 0.27 | 0.79 |
| Openness |  |  |  |  |
| Intercept | -0.09 | 0.25 | -0.36 | 0.72 |
| Male vs. Female | 0.59 | 0.22 | 2.67 | 0.009 |
| Age | -0.05 | 0.03 | -1.34 | 0.18 |
| T allele present | -0.20 | 0.20 | -0.99 | 0.32 |
| Negative Affect |  |  |  |  |
| Intercept | 0.36 | 0.25 | 1.44 | 0.15 |
| Male vs. Female | -0.15 | 0.22 | -0.67 | 0.5 |
| Age | -0.05 | 0.03 | -1.49 | 0.14 |
| T allele present | 0.02 | 0.20 | 0.09 | 0.93 |
| Pro-sociality |  |  |  |  |
| Intercept | 0.67 | 0.23 | 2.92 | 0.004 |
| Male vs. Female | -0.77 | 0.20 | -3.79 | < 0.001 |
| Age | 0.03 | 0.03 | 0.87 | 0.39 |
| T allele present | -0.25 | 0.19 | -1.36 | 0.18 |
| Boldness |  |  |  |  |
| Intercept | -0.26 | 0.25 | -1.03 | 0.30 |
| Male vs. Female | 0.45 | 0.22 | 2.04 | 0.044 |
| Age | 0.00 | 0.03 | 0.00 | > 0.99 |
| T allele present | -0.13 | 0.20 | -0.67 | 0.51 |

*Note*. *N* = 123.
