## Supplementary figures and images for "Personality, Subjective Well-Being, and the Serotonin 1a Receptor Gene in Common Marmosets (*Callithrix jacchus*)"

### Supplementary figures

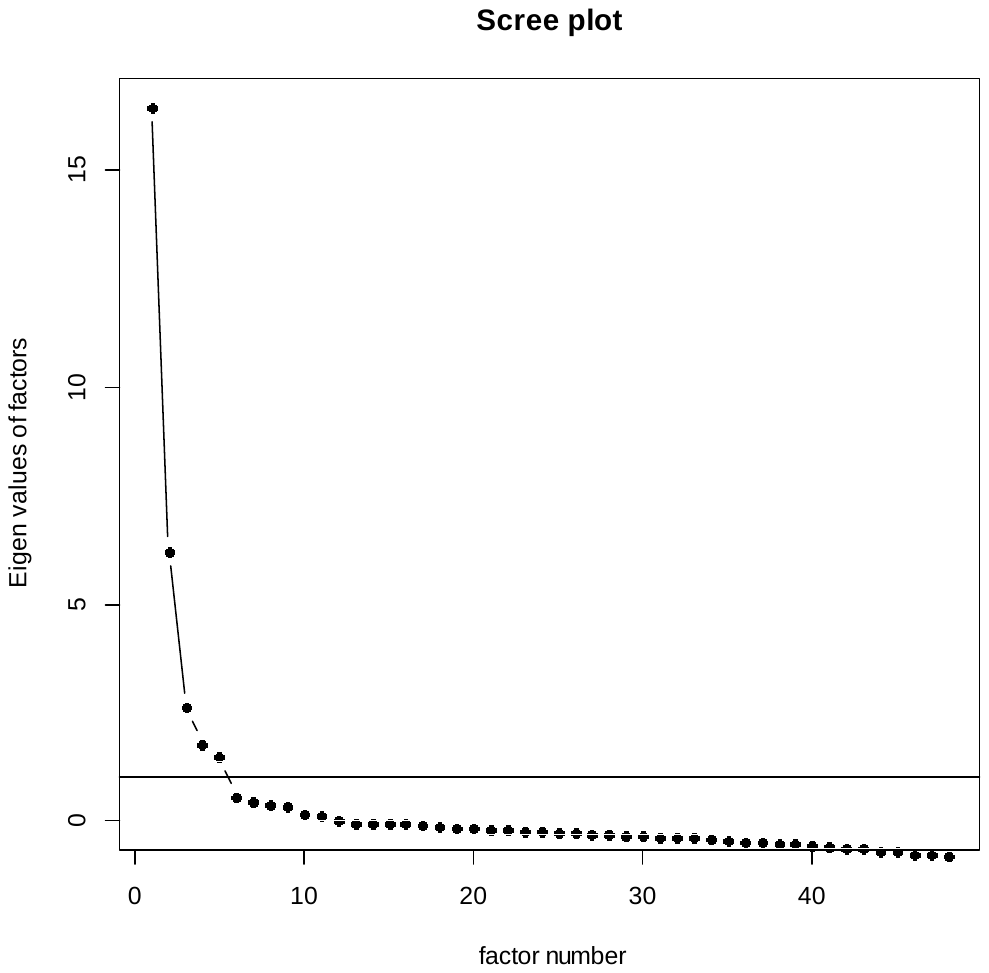

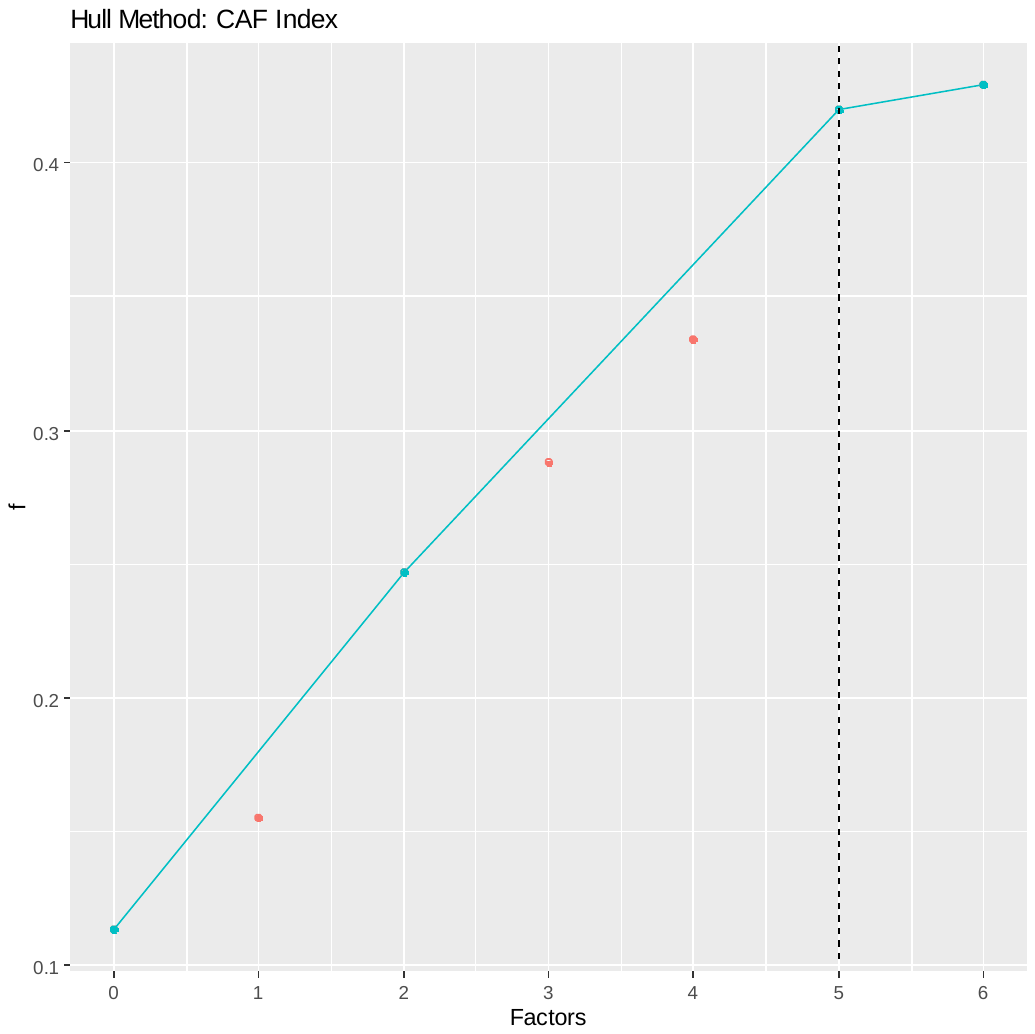

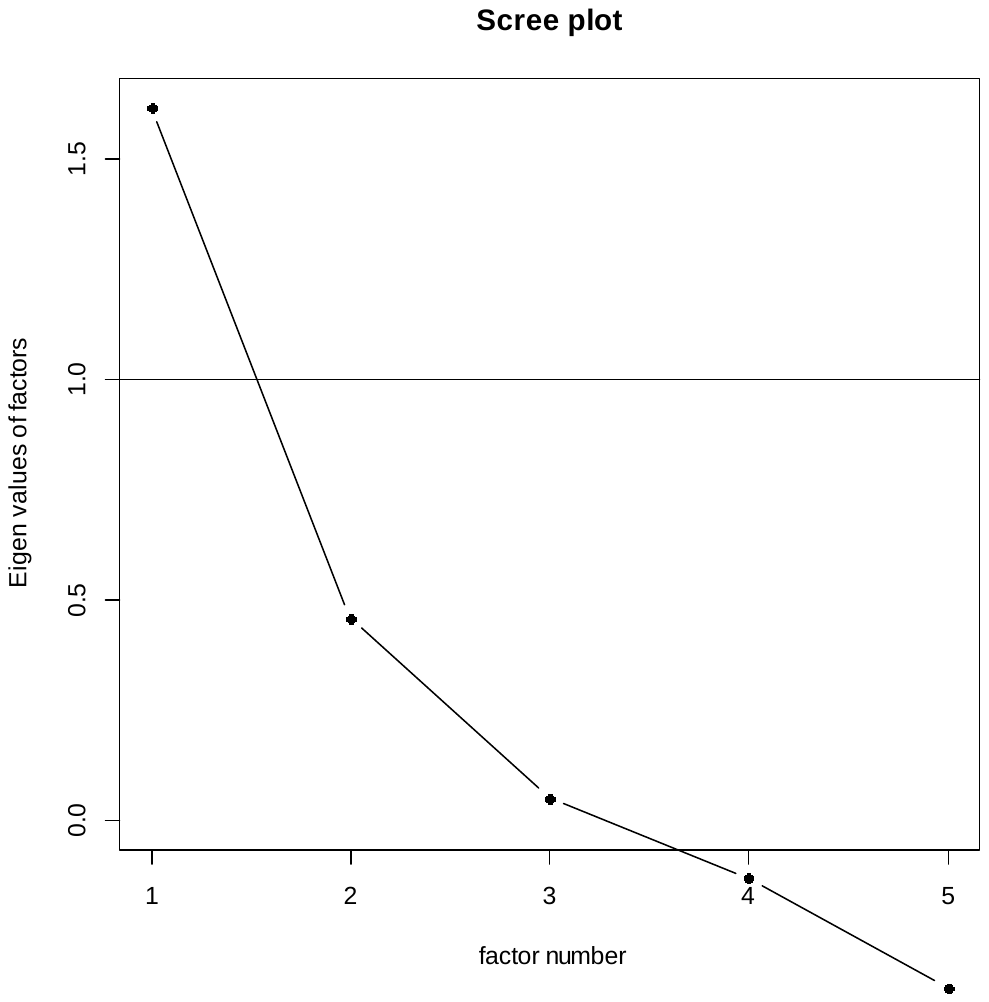

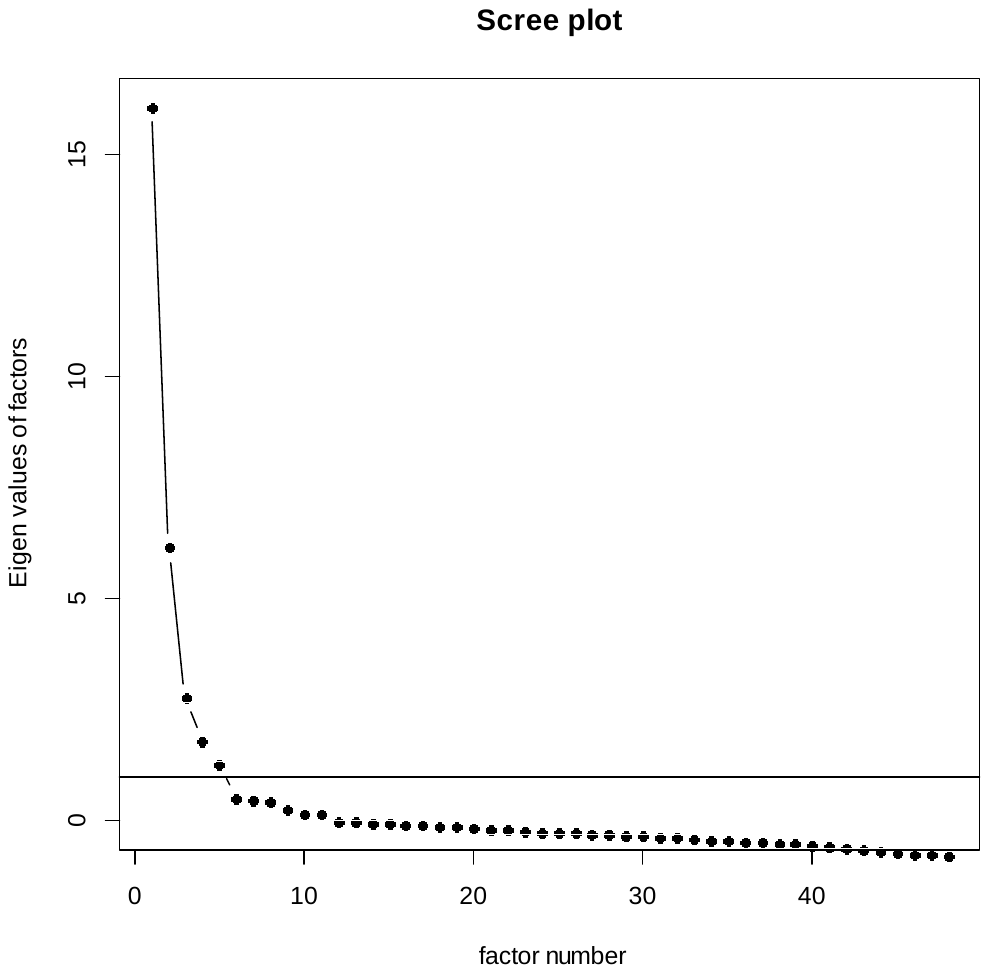

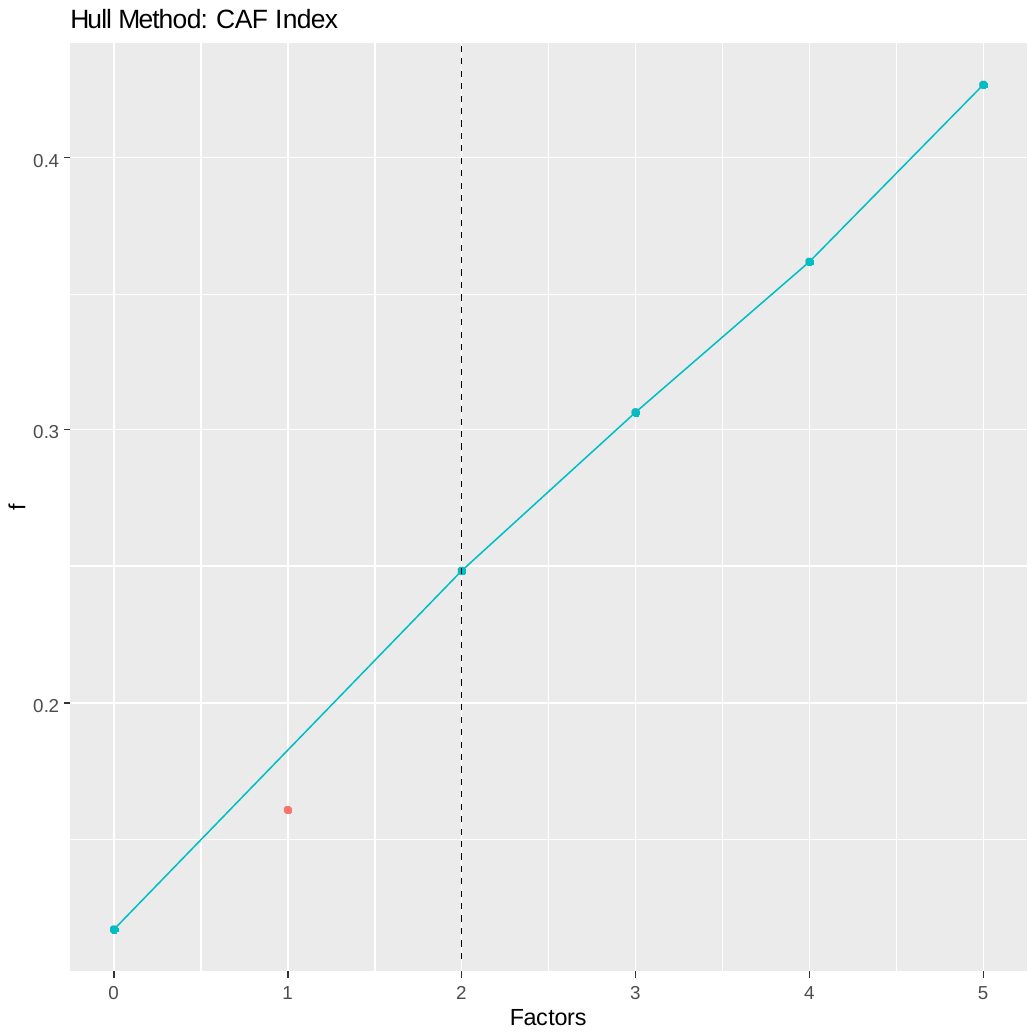

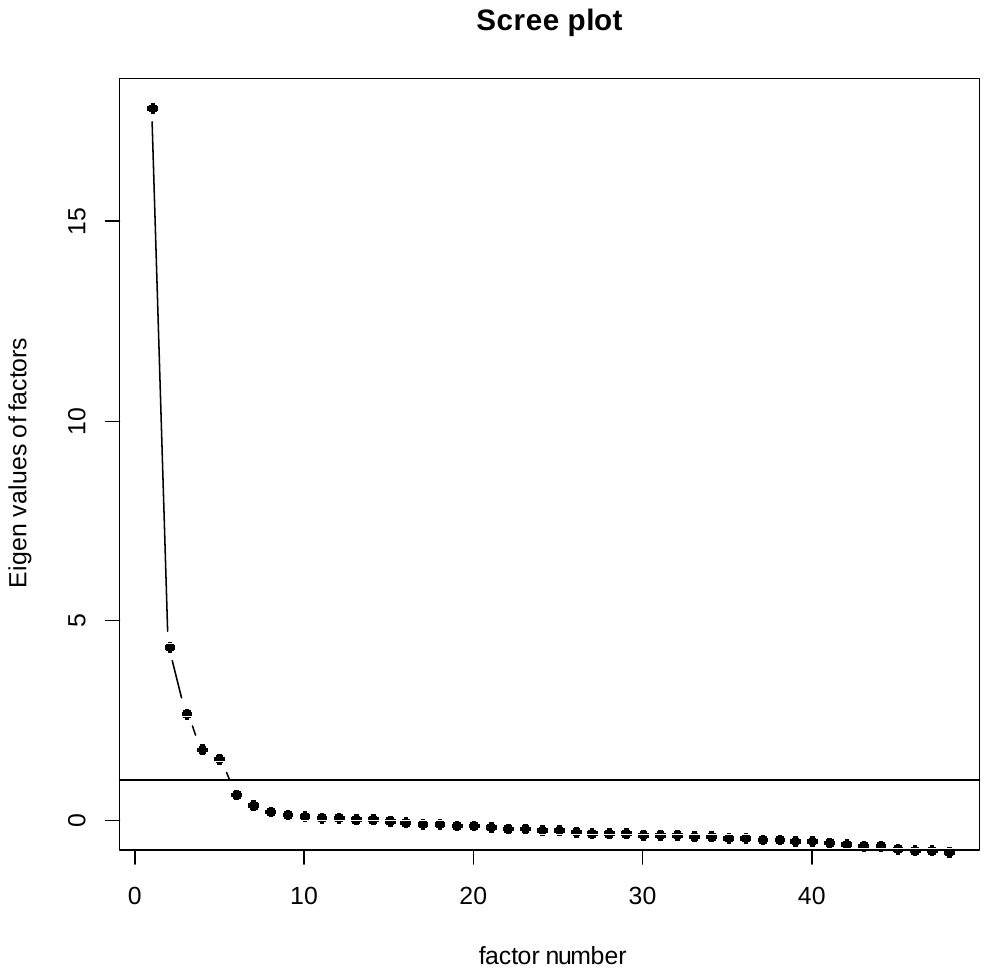

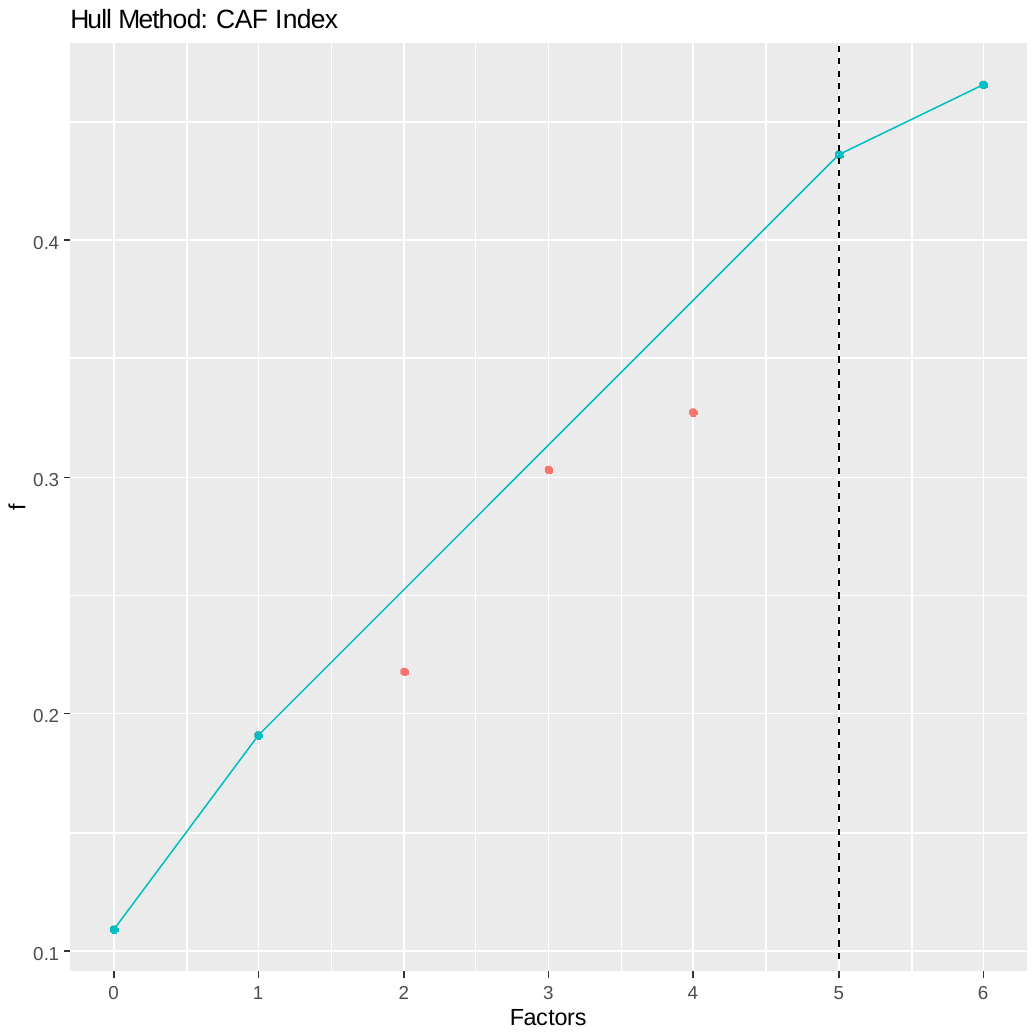

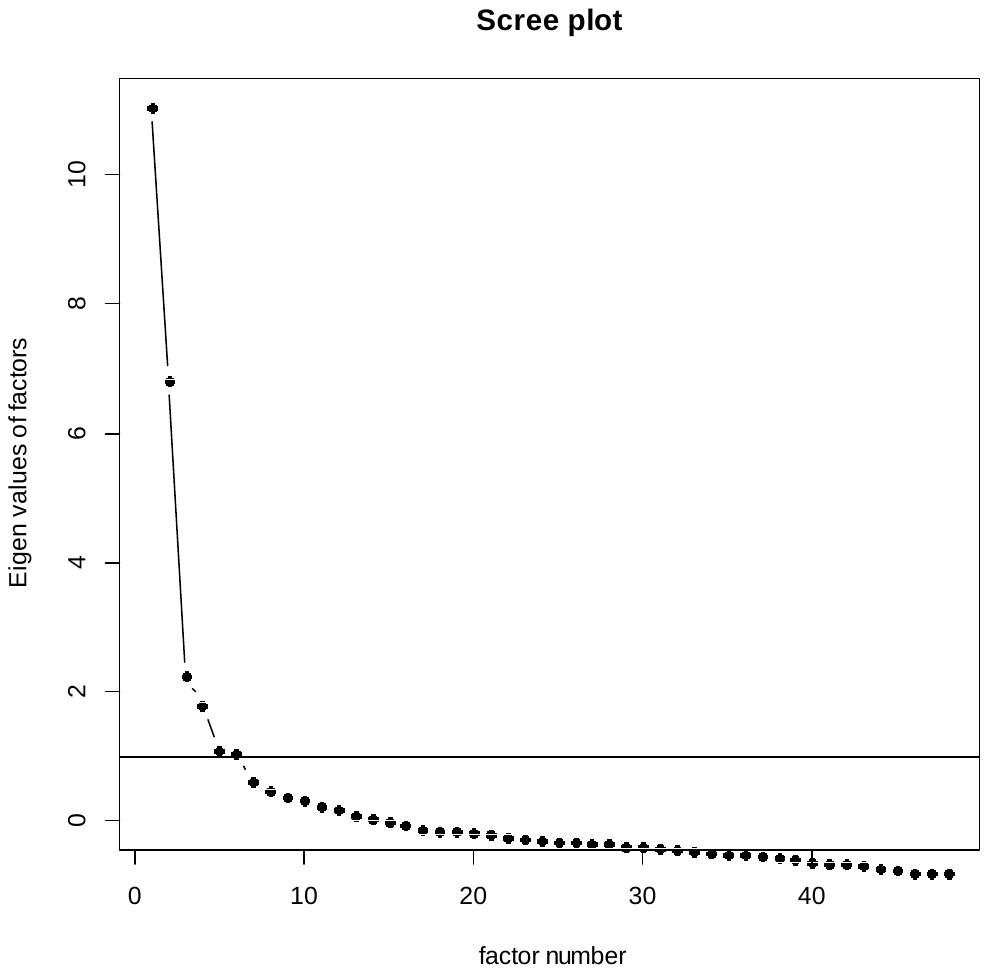

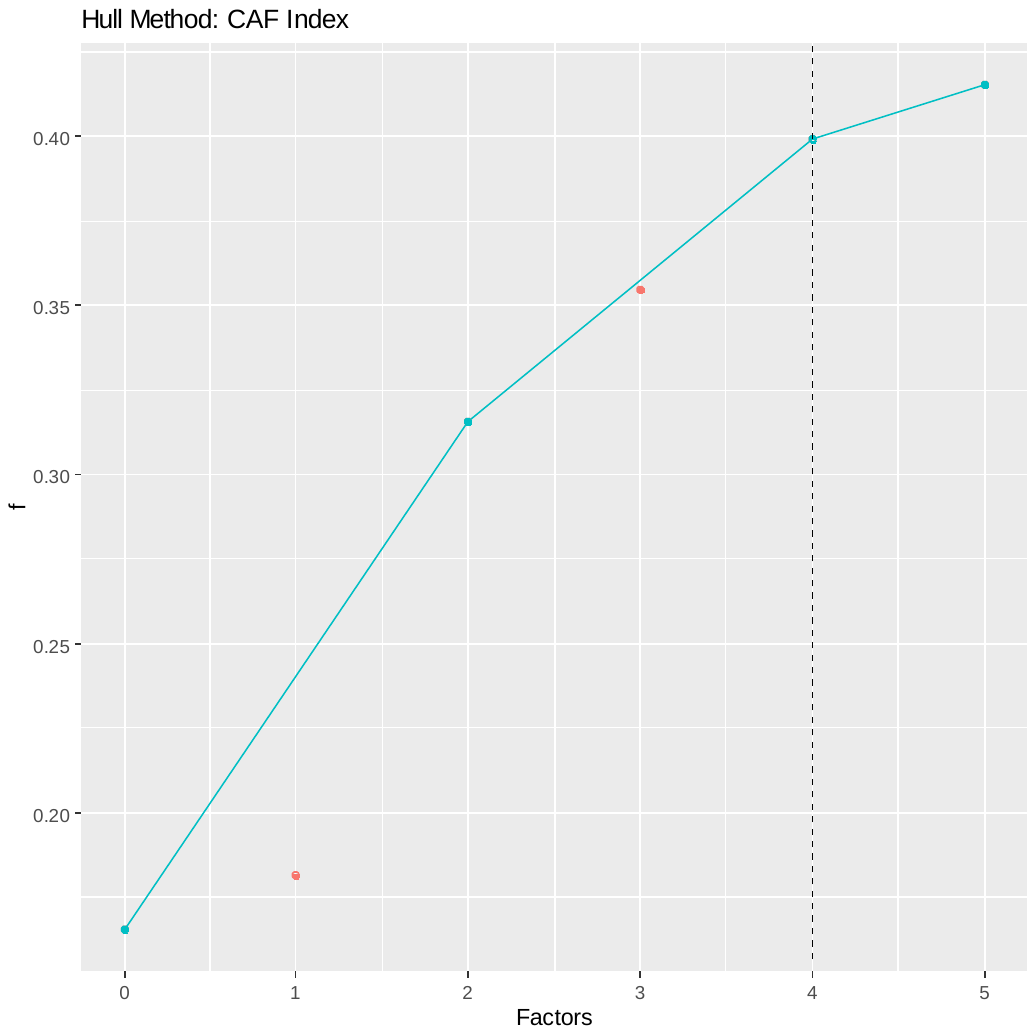

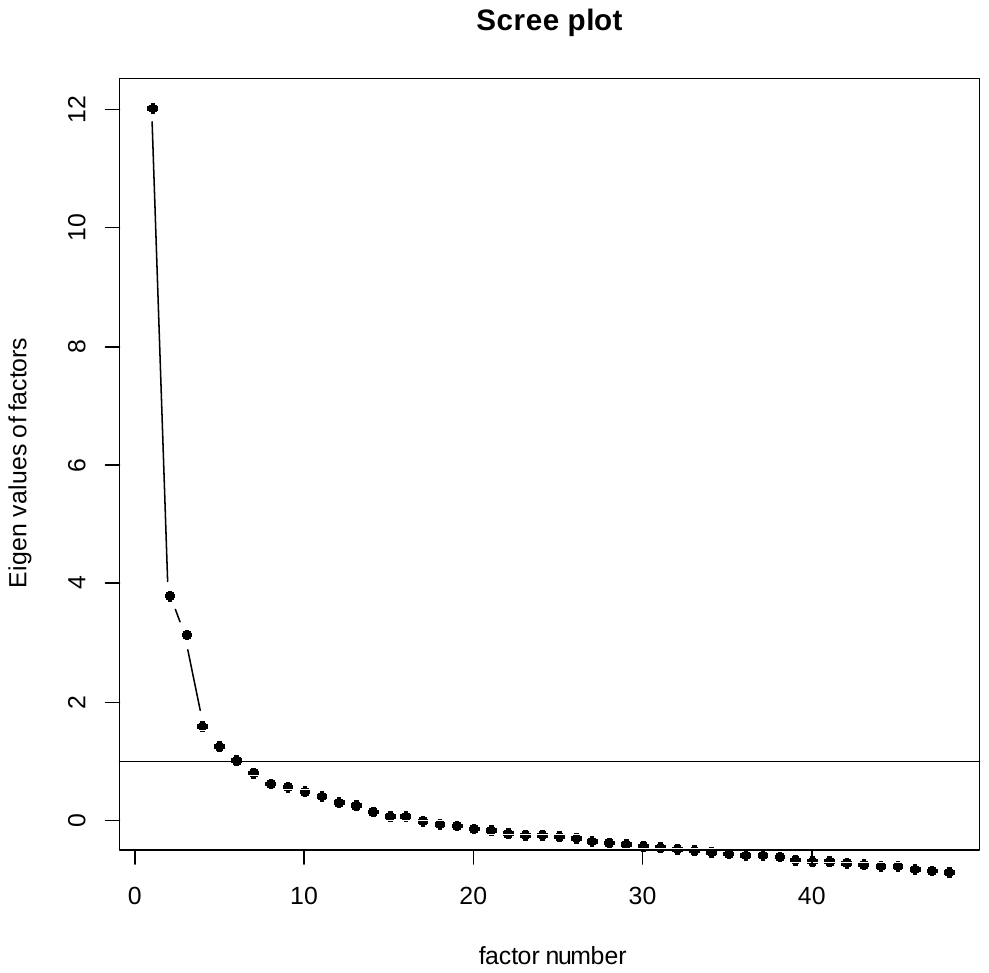

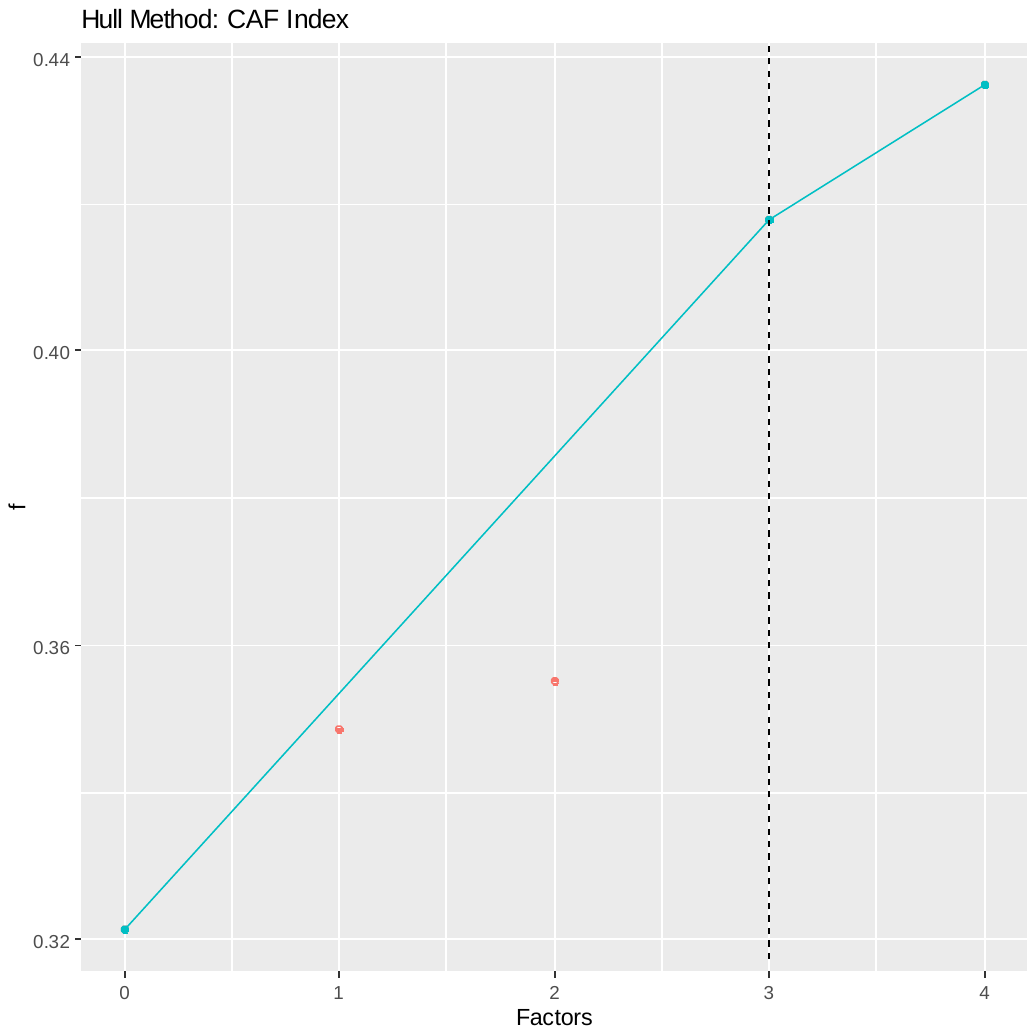

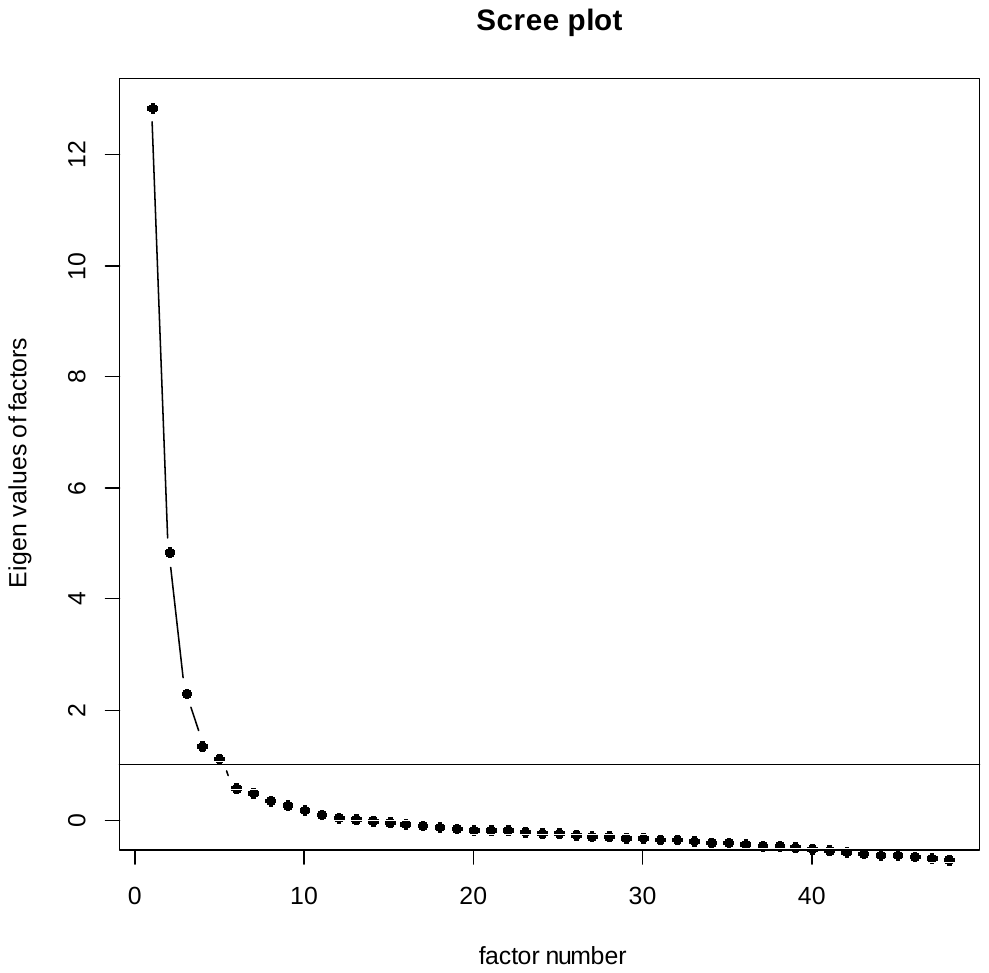

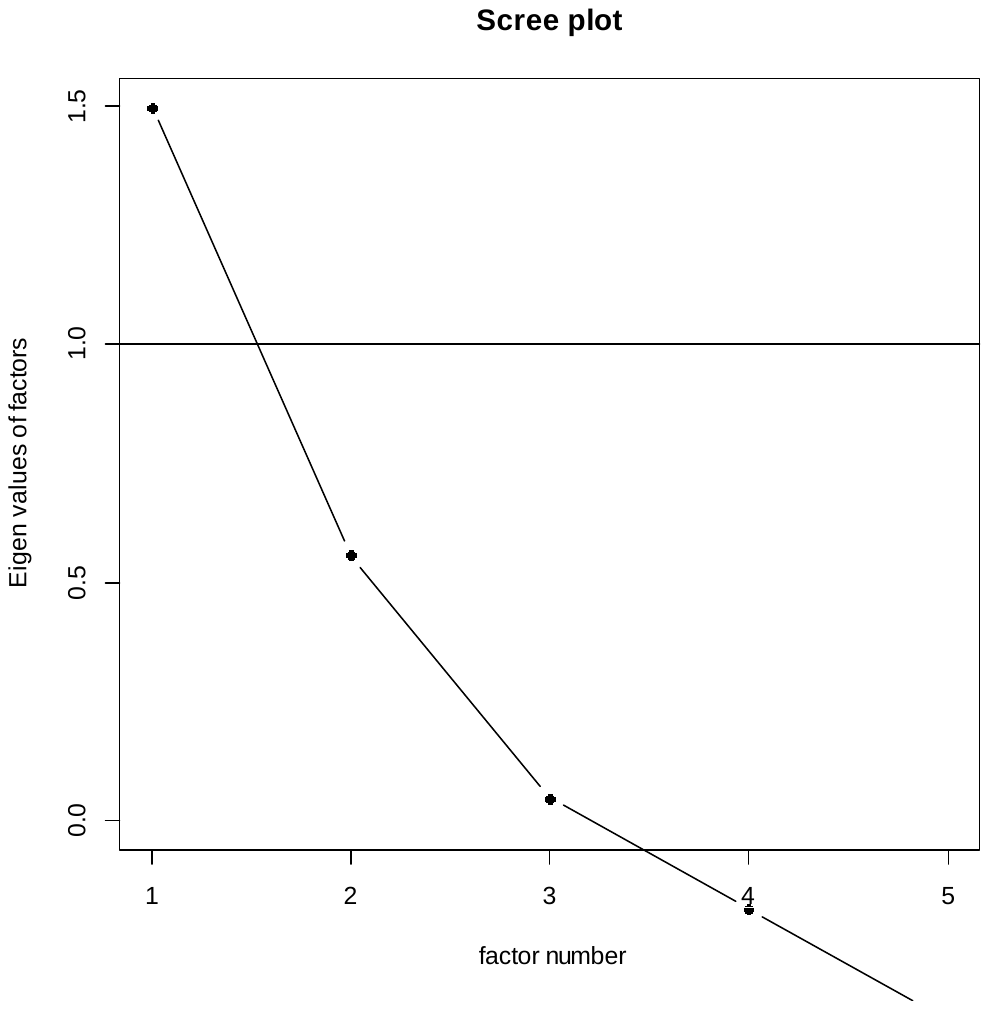

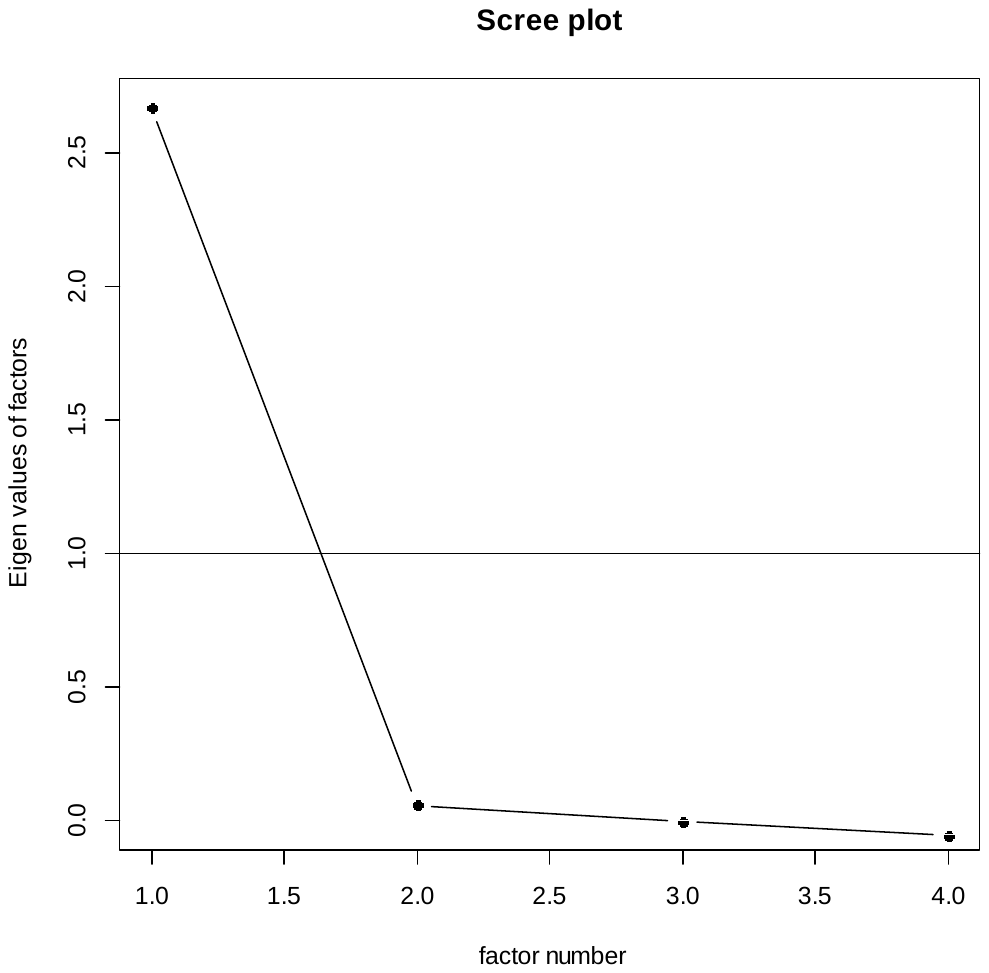
